# Regulated use of alternative Transcription Start Sites controls the production of cytosolic or mitochondrial forms of branched-chain aminotransferase in *Kluyveromyces marxianus*

**DOI:** 10.1101/2022.04.27.489738

**Authors:** Angela Coral-Medina, Darren A. Fenton, Javier Varela, Carole Camarasa, Pavel V. Baranov, John P. Morrissey

**Affiliations:** School of Microbiology, University College Cork, Ireland; SPO, Univ Montpellier, INRAE, Institut Agro, Montpellier, France; School of Biochemistry and Cell Biology, University College Cork, Ireland; Metabolic engineering department, CarboCode GmbH, Konstanz, Germany

## Abstract

Following a whole genome duplication (WGD) event approximately 100 million years ago, the yeast lineage from which the model *Saccharomyces cerevisiae* derives maintained two copies of genes where it was necessary to synthesise proteoforms with different sub-cellular localisation. In contrast, yeasts that did not undergo the WGD event have a single gene that must encode both proteoforms. We adopted an integrated *in silico* and experimental approach to study how this is achieved with *BAT1*, a gene that encodes mitochondrial and cytosolic forms of a branched chain aminotransferase (BCAT) in pre-WGD yeast such as *Kluyveromyces marxianus*. We determined that condition-specific regulation of alternative transcription sites gives rise to mRNA isoforms that differ at the 5’end and that, when decoded, generate a mitochondrial or cytosolic proteoform. Furthermore, targeted mutants lacking specific transcription factors were generated to establish how this differentiation was regulated. As in *S. cerevisiae*, Gcn4 and Leu3 activated expression of the mRNA encoding the mitochondrial proteoform under conditions when branched chain amino acid synthesis was required. Unlike *S. cerevisiae*, however, *K. marxianus* lacked tight regulation of the mRNA encoding the cytosolic proteoform supporting the hypothesis that maintaining paralogous genes in post-WGD yeasts facilitated development of more sophisticated expression control mechanisms.

## INTRODUCTION

Many volatile molecules derived from yeast metabolism have flavour and aroma properties that make significant contributions to fermented beverages and are of increasing interest for the food and cosmetic sectors (Dzialo et al., 2017). Some of the most significant aromas and flavours are derived from the degradation (catabolism) of amino acids in the Ehrlich pathway to produce a range of higher alcohols, acids and esters. These include the aromatic amino acids, phenylalanine and tyrosine, and the branched chain amino acids (BCAA), valine, isoleucine and leucine, and individual amino acids such as threonine and methionine (Hazelwood et al, 2008; Isogai et al., 2022). In each case, the first catabolic step is a transamination to form an α-ketoacid by distinct aromatic amino acid (AAT) or branched chain amino acid aminotransferases (BCAT). The reaction is reversible and, depending on the nutritional requirements of the cell, the pathway can be used for either the degradation or the biosynthesis of amino acids. There is a degree of pathway compartmentalisation with the aminotransferase catabolic reaction taking place in the cytosol and the reverse biosynthetic reaction in the mitochondrion. Although most work on flavour/aroma metabolism has been performed in the baker’s yeast, *Saccharomyces cerevisiae*, the potential for non-traditional yeasts as sources of these metabolites has led to new interest in the study of amino acid metabolism and production of flavours and aromas in non-*Saccharomyces* species (Morrissey et al., 2015).

*S. cerevisiae* often deals with the challenge of producing cytosolic and mitochondrial variants of the same enzyme (proteoforms) by possessing two copies of the relevant gene, one of which encodes a protein with a mitochondrial targeting signal (MTS). Examples of this include *BAT1* and *BAT2,* or *AAT1* and *AAT2,* encoding BCAA and aspartate aminotransferases, respectively. These are paralogous genes (Kellis et al., 2004) that arose from the whole genome duplication event that occurred approximately 100 million years ago when ancestral yeasts from the *Kluyveromyces*, *Lachancea* and *Eremothecium* (KLE) and *Zygosaccharomyces / Torulaspora* (ZT) clades hybridised to form a new lineage (Conant and Wolf, 2008; Marcet-Houben and Gabaldón, 2015; Wolfe, 2015). The presence of paralogous genes post-WGD enabled functional differentiation in some cases, for example, differential gene expression, alteration of kinetic properties, oligomeric organization or sub-cellular localisation of expressed proteins (González et al., 2017; Quezada et al., 2008). In contrast, so-called “pre-WGD” yeast such as those in the KLE clade, retain the ancestral state of a single gene that must be responsible for both functions of the orthologous *S. cerevisiae* genes. Establishing how this can be achieved is important for understanding the metabolism and physiology of non-*Saccharomyces* yeasts. It also contributes to our understanding of some of the evolutionary processes that took place post-WGD leading to the specialisation and niche adaptation of *S. cerevisiae*.

*BAT1,* encoding the mitochondrial BCAT, and *BAT2,* the cytosolic one, are among the best studied aminotransferases in *S. cerevisiae*. Bat1 and Bat2 have 77% amino acid identity and can each perform the forward or reverse reaction but the Bat1 proteoform is localised in the mitochondrial matrix due to the presence of a mitochondrial targeting signal (MTS) at the amino terminus (Kispal et al., 1996). The different sub-cellular localisation allows functional specialisation of Bat1 for BCAA synthesis in the mitochondrion and Bat2 for BCAA degradation in the cytosol (Colon et al. 2011). *BAT1* and *BAT2* show different patterns of expression depending on the environmental nitrogen conditions and the related physiological requirements of the cell (González et al., 2017). The nitrogen-source control of *BAT1/BAT2* expression is tight, with both genes being subject to the activity of transcriptional regulators. In *S. cerevisiae* grown in biosynthetic conditions (for example using NH_4_ or glutamine as a sole nitrogen source), the transcription factors (TFs) Gcn4 and Leu3 directly bind to the *BAT1* promoter activating expression. The activating function of Leu3 requires that it is bound by *α*-isopropylmalate (*α*-iPM), an intermediate of the leucine biosynthetic pathway – in the absence of (*α*-iPM), Leu3 represses transcription. There is no *BAT2* expression under these conditions and nucleosomes occlude much of the promoter. In contrast, under catabolic conditions (BCAA as a nitrogen source), *BAT1* is repressed and *BAT2* is activated. The activation of *BAT2* involves the nitrogen catabolite repression (NCR) activator Gln3, which translocates to the nucleus under these conditions and directly binds the *BAT2* promoter, driving expression. A second TF, Put3, also binds the *BAT2* promoter and has an activating effect. Put3 plays a second role in this system as it represses *LEU1*, which is required for *α*- iPM synthesis. The effect of this is that the Leu3 bound to the *BAT1* promoter is not bound by *α*-iPM and so represses *BAT1* expression. As these are amino acid-replete conditions, Gcn4 levels are low so this activator is absent and thus cannot overcome the repressive effect of Leu3.

Previous studies of the BCAT genes in pre-WGD yeasts, *Kluyveromyces lactis* and *Lachancea kluyveri* reported that two proteoforms with opposing functions are encoded by a single gene *BAT1* (Colón et al., 2011; Montalvo-Arredondo et al., 2015). Those studies proposed that the biosynthetic and catabolic roles of the ancestral Bat1 have been partitioned in *BAT1* and *BAT2* through differential expression and subcellular re-localisation, improving regulation of BCAA metabolism in *S. cerevisiae* (Montalvo-Arredondo et al., 2015). A number of questions remained unresolved from the previous studies. First, how does the single *BAT1* gene give rise to both cytosolic and mitochondrial proteoforms? Second, how is that process regulated and controlled? Addressing this leads to a further question as to whether the nitrogen source-dependent transcriptional regulation of *BAT1* and *BAT2* that is seen in *S. cerevisiae* is derived from ancestral regulation or was newly-acquired in post-WGD yeasts? We chose to address these questions in *Kluyveromyces marxianus* for three main reasons. First, a recently published multi-omics based annotation of *K. marxianus* provides a level of genome-wide resolution on gene structure and expression and gives the raw data to investigate the *BAT1* locus in detail (Fenton et al., 2022a, 2022b). Second, the CRISPR-based genome engineering tools developed for *K. marxianus* enable rapid and precise genome editing thereby facilitating strain engineering to address hypotheses that are generated (Rajkumar and Morrissey, 2022). And finally, because *K. marxianus* is emerging as one of the most important new yeasts for biotechnology applications, there is an imperative to improve our knowledge of its genetics and physiology (Fonseca et al., 2008; Homayouni-Rad et al., 2020; Karim et al., 2020; Leonel et al., 2021; Morrissey et al., 2015).

This study set out with the aim of understanding how *K. marxianus* manages to produce each Bat1 proteoform depending on the metabolic needs. Initially, the structure of the *BAT1* locus was examined in the GWIPS-Viz genome browser (Michel et al., 2017) to identify features such as transcriptional start sites (TSS) (TSS), mRNA translation and ORF architecture to examine potential alternative translation start sites. Next, using protein fusions, we demonstrated the presence of an MTS at the N-terminus of Bat1. Expression analysis confirmed that *BAT1* uses alternative TSS to produce the two differentially localised Bat1 proteoforms, with the choice of TSS determined by the nitrogen status of the cells. To investigate this further, we individually inactivated eight transcript factors (TFs) and found that several of those involved in regulation of *BAT1* and *BAT2* in *S. cerevisiae* also play a role in *BAT1* regulation in *K. marxianus*. The main similarity was the positive effect of Gcn4 and Leu3 on expression of the mitochondrial form of BCAT. There was a divergence with the cytosolic form, however, with the data indicating the paralogue *BAT2* acquired positive regulation via Gln3 and Put3 following the whole genome duplication.

## MATERIALS AND METHODS

### Strains and culture conditions

All *K. marxianus* strains used in this study are derived from *K. marxianus* NBRC1777 and are listed in Table 1. Strains were routinely cultivated in YPD broth (2% glucose, 1% yeast extract, 2% peptone) at 28°C. For experiments with specific nitrogen sources, a minimal media (MM) based on the standard Verduyn medium was used (glucose 20 gL^-1^, KH_2_PO_4_ 3 gL^-1^, MgSO_4_.7H_2_O 0.5 gL^-1^, vitamin mix, and trace elements mix) (Verduyn et al., 1992). This was supplemented with individual nitrogen sources as follows: ammonium sulfate 5 gL^-1^, glutamine 5.5 gL^-1^, valine 8.8 gL^-1^, isoleucine 9.9 gL^-1^, leucine 9.9 gL^-1^, or a mixture of BCAAs termed VIL (valine 150 mgL-1, isoleucine 30 mgL-1, leucine 100 mgL-1). For tests on different N-sources, 10 mL cultures were pre-grown on YPD for 16h, harvested by centrifugation, and resuspended in Yeast Nitrogen Base (YNB) (glucose 20 gL^-1^) media without amino acids and ammonium sulphate. Incubation was continued for further 4h to deplete cellular nitrogen reserves and the cells were harvested again, washed with 0.9% NaCl saline solution, before resuspension in the desired growth medium at an initial A_600_ of ∼0.05. Subsequent growth tests were performed either in flasks or microtitre plates. When required for selection and plasmid maintenance, G418 was used at 200 ng/μL.

**Table 1.** Yeast strains used in this study. Coordinates of the mutations in the *K. marxianus* NBRC1777 genome are shown. The deleted nucleotides are represented by Δ in cases where several nucleotides were deleted.

| Strain name | Genotype of mutants | Source |
| --- | --- | --- |
| *Kluyveromyces marxianus* NBRC1777 | Wild type | NITE Biological Research Centre |
| *K. marxianus* NBRC1777 Δlig4 | Lig4-1 Frameshift | (Rajkumar et al., 2019) |
| *K. marxianus* NBRC1777 Δgat1 | CHR VI Δ635,534 – 636,959 | This study |
| *K. marxianus* NBRC1777 Δgcn4 | CHR V Δ868,076 – 868,697 | This study |
| *K. marxianus* NBRC1777 Δgln3 | CHR III Δ1,456,475 – 1,457,877 | This study |
| *K. marxianus* NBRC1777 Δhap2 | CHR III Δ1,550,967 – 1,551,414 | This study |
| *K. marxianus* NBRC1777 Δmot3 | CHR VI Δ700,804 – 701,788 | This study |
| *K. marxianus* NBRC1777 Δnrg1 | CHR IV Δ283,460 – 284,113 | This study |
| *K. marxianus* NBRC1777 Δput3 | CHR VI Δ343,875 – 345,730 | This study |
| *K. marxianus* NBRC1777 Δleu3 | CHR VI Δ946,030 – 947,790 | This study |
| *K. marxianus* NBRC1777 Δbat1 | CHR I Δ1,068,093 – 1,069,223 | This study |
| *K. marxianus* NBRC1777 ΔNTE-Bat1 | CHR I Δ1,068,052 – 1,068,127 | This study |

Batch growth was performed in 100 mL flasks containing 25 mL of MM supplemented with the appropriate nitrogen source. The cultures were inoculated at initial A_600_ ∼0.05. Experiments were performed in triplicate. For the microplate fermentations the A_600_ of the starter cultures was set to 0.5 to then inoculate fresh media at ∼0.05 initial A_600_; the total volume in each well was 200 μL. This experiment was done with a minimum of four biological replicates of each strain in each media (ammonium sulfate, VIL, glutamine, valine, isoleucine, leucine, ammonium+VIL), using one blank per media. The 96 flat wells microplate was incubated in the microplate reader CLARIOStar®^Plus^ (BMG LABTECH, Germany). The absorbance at 600 nm was measured for 60 cycles of 1 hour (24 flashes/cycle) with continual double orbital shaking (400 rpm) between measurements.

### Construction of deletion mutants

CRISPR-Cas9 was used to construct *K. marxianus* deletion mutants by introduction of a repair fragment as previously described (Rajkumar and Morrissey, 2022). That protocol describes the detailed construction of plasmids to create a double stranded break (DSB) in a target gene and a repair fragment that integrates at this site to introduce a deletion that inactivates the gene. Each plasmid along with the repair fragment that introduced a deletion of between 500 and 1500 bp (depending on the gene) was transformed into *K. marxianus* NBRC1777 *Δlig4* (Table 1) using the LiAc/SS carrier DNA/PEG method (Gietz and Schiestl, 2007). Transformant strains were checked by PCR with diagnostic primers and DNA sequencing (Eurofins Genomics, Germany) to confirm the deletion in the target locus. The primers used for plasmid construction, verification and amplifications are listed in supplementary table 1. Further details of each mutant are provided in supplementary material 1.

### Gene expression analysis

For gene expression analysis, batch growth was performed in 100 mL flasks containing 25 mL of MM supplemented with the appropriate nitrogen source. The cultures were inoculated at initial A_600_ ∼0.05 and the cells were collected by centrifugation at exponential phase when the A_600_ reached approximately 0.8. The pellets were immediately snap-frozen in liquid nitrogen and RNA was extracted with the Trizol method (Chomczynski and Sacchi, 1987) using glass beads as described by Tesnière et al., (2021). RNA was purified with the RNeasy Mini Kit (QIAGEN) following the RNA clean-up protocol, using DNase I for DNA digestion to remove the contaminating DNA present in the samples. The purified RNA quality was evaluated through capillary electrophoresis in the Bioanalyzer 2100 with an RNA 6000 Nano LabChip Kit (Agilent Technologies, Santa Clara, CA, USA) and quantified using a Nanodrop spectrophotometer. All the samples were diluted to the same RNA concentration at 160 ng/μL. cDNA synthesis was performed using the SuperScript III Reverse Transcriptase Kit (Invitrogen), with 100 ng/μL of random primers and 5 μL of RNA for a total reaction volume of 20 μL. The cDNA synthesis was also done for all samples without the Reverse Transcriptase III (-RT) to use as control. After this step, the cDNA samples were used as template for Real-Time qPCR with technical triplicates. The qPCR reaction of 25 μL final volume, contained 5 μL of template cDNA, 0.3 pmol/μL of each primer and 12.5 μL 2X Power SYBR Green I PCR Master Mix (Applied Biosystems, Warrington, UK). The thermocycling conditions were as follows: an initial enzyme activation of 2 minutes at 50°C and 10 minutes at 95°C, followed by 40 cycles of denaturation for 15 seconds at 95°C, annealing and extension for 1 minute at 60°C, with a final melt gradient starting from 15 seconds at 95°C, 30 seconds at 60°C and finally 15 seconds at 95°C. The real-time qPCRs were carried out in a 7300 Fast Real Time PCR System (Applied Biosystems). Primer specificity was confirmed by analysing dissociation curves of the PCR amplification products. The primers used for the amplification of BAT1-Long, BAT1-Total, and ACT1 mRNA are listed in supplementary table 1. Standard curves were done to calculate the primer efficiency of each primer set using genomic DNA of *K. marxianus NBRC1777* extracted with (LiOAc)-SDS method adapted from (Lõoke et al., 2011). The data was normalized against *ACT1* used as control. The relative gene expression was calculated with Pfaffl mathematical model that determines the relative expression ratio (R) of the target genes based on the primer efficiency (E) and the Ct deviation of the genes of interest versus the *ACT1* control (Pfaffl, 2001). The results were plotted and significant differences between the mutants and conditions were calculated using the two-way analysis of variance (ANOVA) in GraphPad Prism. The differences were considered statistically significant when the p-values were <0.001.

### Heterologous expression of fusion proteins

Three plasmids were constructed for the protein localisation experiments: pBAT1-L, pBAT1- S, pNatProm-BAT1 (Supplementary figure 1). For the pBAT1-L and pBAT1-S construction, the BAT1 short and long open reading frame fragments were amplified by PCR from *K. marxianus NBRC1777* gDNA using Q5 polymerase (New England Biolabs (NEB) Inc., MA, USA). The fragments were then cloned into pMTU-DO-G418-C5 (Table 2) with Golden Gate Assembly, together with the GFP open reading frame part and the *TEF1* promoter (Rajkumar and Morrissey, 2020). The link between *BAT1* and GFP was 9 nucleotides long and encoded phenylalanine, tryptophan and glycine amino acids designed to form an in-frame fusion. For the construction of pNatProm-BAT1, the fragments used were the *BAT1* native promoter that consists of 1000 bp upstream of the initiator AUG including the N-terminal extension (NTE), obtained by PCR from *K. marxianus* gDNA, and the GFP fragment with the *INU* terminator obtained by PCR from the pBAT1-S plasmid previously constructed. These fragments were cloned with the pMTU-DO-G418-C5 backbone using Gibson Assembly Master Mix (NEB) following the manufacturer’s instructions. The assemblies were transformed into *E. coli* and then plated in LB medium supplemented with kanamycin 50 ng/μL. The plasmids were extracted and verified with HindIII, BstEII, NruI enzymatic digestion (NEB) and DNA sequencing (Eurofins Genomics, Germany). The resulting plasmids were transformed into *K. marxianus* NBRC1777 using the LiAC/SS carrier DNA/PEG procedure. The transformed strains were plated onto YPD agar supplemented with G418 200 ng/μL.

**Figure 1.**
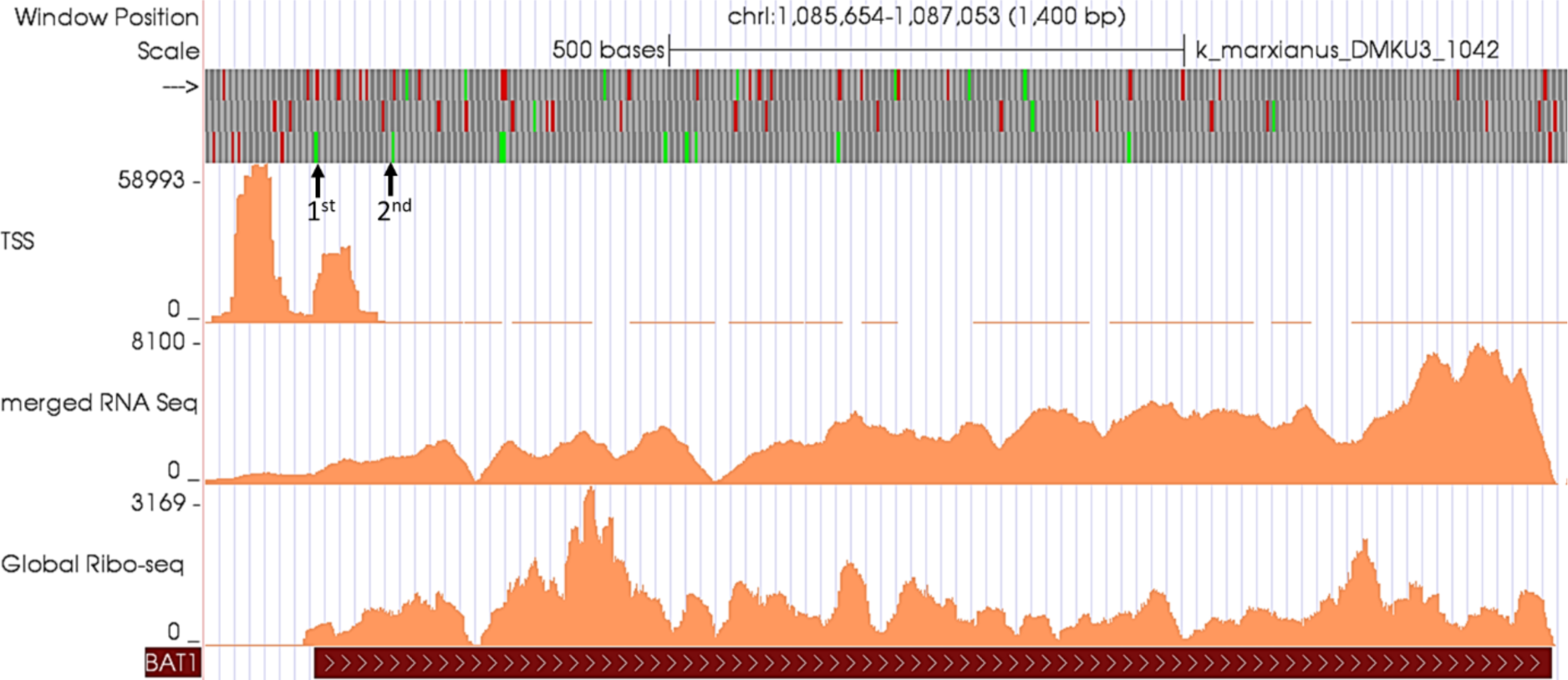
*K. marxianus BAT1* locus visualised on the GWIPS-Viz Genome Browser. The annotation track (red bar) at the bottom represents the coding region of *BAT1*. Coverage tracks of Ribosome profiling (Ribo-Seq), RNA Sequencing (RNA-Seq) and Transcription Start Site Sequencing (TSS-Seq) corresponding to the *BAT1* locus of *K. marxianus* are depicted. The top track represents ORF architecture including start codons in green and stop codons in red. The two black arrows indicate the positions of the first and the second in-frame start codons of *BAT1*. There are two clear TSS at the TSS-Seq track that are located upstream of each start codon.

**Table 2.** Plasmids used in this study

| Plasmid | Source |
| --- | --- |
| pUDP002-HH-BsaI | (Varela et al., 2019) |
| pMTU-DO-G418-C5 | (Rajkumar and Morrissey, 2020) |
| pBAT1-L | This study |
| pBAT1-S | This study |
| pNatProm-BAT1 | This study |
| pCRISPR-KmΔgat1 | This study |
| pCRISPR-KmΔgcn4 | This study |
| pCRISPR-KmΔgln3 | This study |
| pCRISPR-KmΔhap2 | This study |
| pCRISPR-KmΔmot3 | This study |
| pCRISPR-KmΔnrg1 | This study |
| pCRISPR-KmΔput3 | This study |
| pCRISPR-KmΔNTEBAT1 | This study |
| pCRISPR-KmΔbat1 | This study |

### Fluorescence microscopy

Cultures of *K. marxianus* NBRC1777 carrying the pBAT1-L, pBAT1-S, pNatProm-BAT1 plasmids were grown at 28°C in 25 mL of MM with either ammonium or VIL as the nitrogen source at the mentioned concentrations. Cells carrying pNatProm-BAT1 plasmid were additionally grown in YPD in the same conditions. The cells were collected at exponential phase by centrifugation and resuspended in pre-warmed staining solution that consist of MitoTracker Red CMXRos (Invitrogen) at 200 nM. The suspensions were incubated for 45 minutes in the same growth medium as above and then washed with fresh pre-warmed 0.9% NaCl saline solution. The fixation was done in pre-warmed growth medium containing 3.7% formaldehyde at 37°C for 15 minutes. Antifade reagent Concanavalin-A (Invitrogen) at 0.5 mg/mL was used as mounting media for visualization. For fluorescence detection, GFP and MitoTracker Red were excited by 488 nm argon and 543 nm helium/neon lasers respectively. All images were captured using an Olympus Fluoview 1000 Confocal Laser Scanning Microscope (Olympus Corporation, Japan) with Fluoview software, using a 100x apochromatic objective with immersion oil.

### Bioinformatic and statistical analysis

Expression of genes at specific genetic loci (BAT1, ALT1, BAP3) was analysed in the *K. marxianus* GWIPS-viz database (Fenton et al., 2022a). The prediction of the presence of a mitochondrial targeting signal (MTS) in Bat1p and the corresponding cleavage site (CS), was done with TargetP 2.0 software (Armenteros et al., 2019). The analysis of the growth kinetics data was performed using R Studio software, version 1.3.1093 (RStudio Team., 2020) with GrowthCurver package (Sprouffske and Wagner, 2016). T graphs and statistical tests were performed using GraphPad Prism version 8.0.2 for Windows (GraphPad Software, San Diego, California, USA). The *K. marxianus BAT1* promoter analyses was performed using the web based software YetFasco (De Boer and Hughes, 2012) and the Yeastract database (Monteiro et al., 2020) to search for putative TF-binding sites.

### Data Availability

Strains and plasmids are available on request. All databases uses in this study are freely accessible: GWIPS-viz genome browser (https://riboseq.org/); Yeastract (http://www.yeastract.com/); YetFasco (http://yetfasco.ccbr.utoronto.ca/). All software listed in the relevant sections is freely available and described in the cited publications. No new code was written for this study. The authors affirm that all data necessary for confirming the conclusions of the article are present within the article, figures, and tables, including the supplemental material provided.

## RESULTS

### K. marxianus BAT1 has two transcription start sites (TSS)

To investigate the mechanism by which mitochondrial and cytosolic isoforms of Bat1 could arise from a single gene in *K. marxianus*, we analysed the *BAT1* locus in the recently published *K. marxianus* GWIPS-viz database (Fenton et al., 2022a). It was possible to identify transcription start sites (TSS-seq), the transcribed region (RNA-Seq) and the translated region (Ribo-Seq) (Figure 1). The two tall histogram peaks in the TSS-seq panel indicate discrete transcription start sites and the RNA-Seq reads show transcription across all this region. There are two possible in-frame translation initiation codons and, from the ribosome profiling data, it is clear that translation can initiate from the first AUG codon. Since the second identified transcription start is downstream of the first AUG codon, a transcript arising from use of this TSS could only initiate translation from the second putative AUG initiation codon. Taking these data together, we hypothesised that two BAT1 mRNA isoforms are produced in *K. marxianus* through the use of alternative TSS (aTSS), each of which may be translated from a different AUG start codon. This would give rise to two different proteoforms that we refer to as Bat1-L (long) and Bat1-S (short). To test if the two versions of Bat1p are predicted to be targeted to different organelles, the subcellular localisation of the two isoforms was analysed using the in silico TargetP 2.0 tool. The program predicted the presence of a MTS in the long isoform of *BAT1*, while no signal peptide was detected in the short isoform (Figure 2). This is in agreement with the prediction of an MTS in this protein by Fenton et al., (2022b). Notwithstanding the presence of two transcriptional starts, an alternative explanation for two isoforms would be leaky scanning, which could lead to bypass of the first AUG. To assess whether this was likely, we examined the context of both AUG start codons (Figure 2). In fact, both have good Kozak consensus, largely excluding the leaky scanning hypothesis. In combination, these data support a model whereby use of alternative transcription start sites, leads to mRNA isoforms that, when translated, give rise to a longer form of Bat1 (Bat1-L) that is localised to the mitochondrion, or a shorter from of the protein that remains cytosolic (Bat1-S).

**Figure 2.**
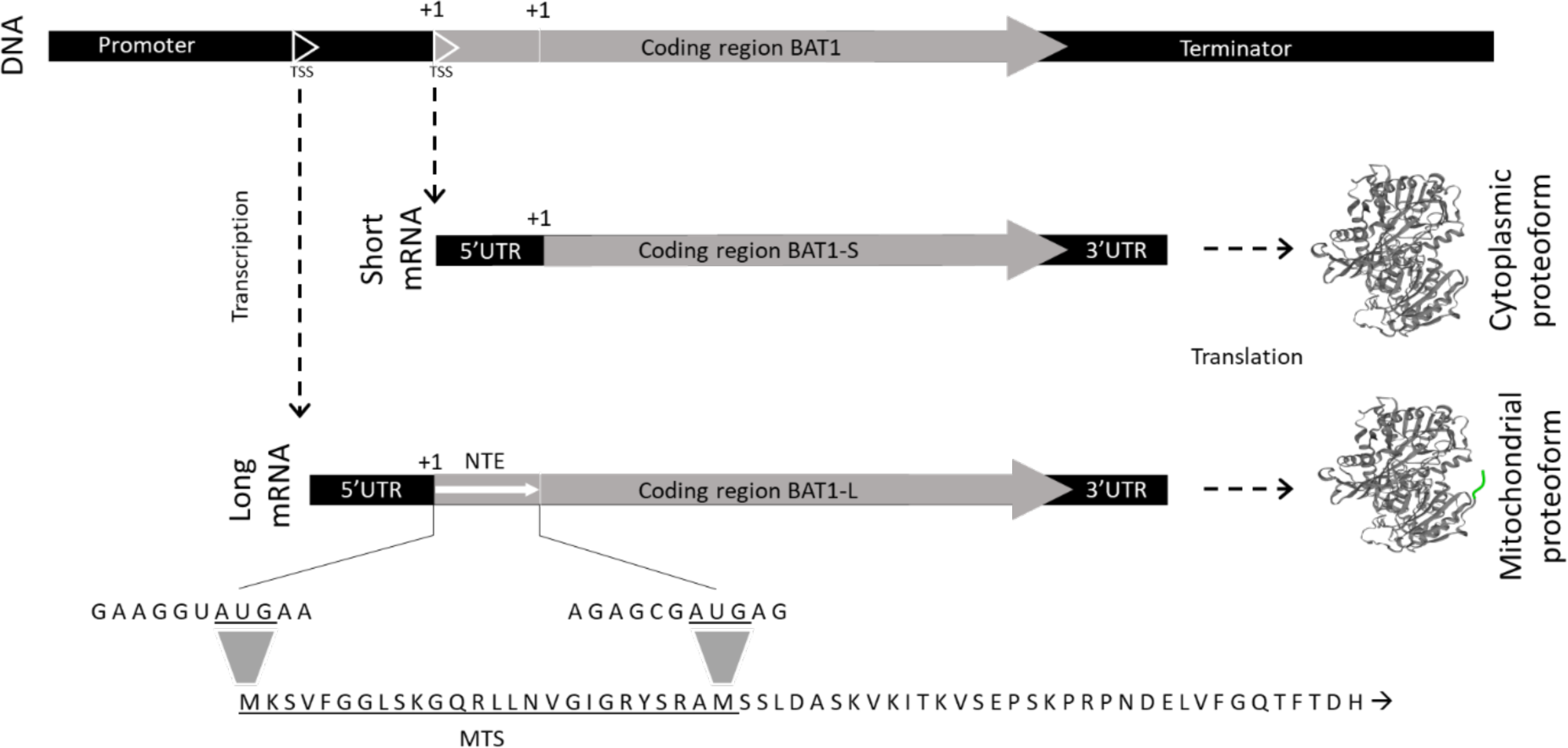
Diagrammatic representation of the production of two Bat1 proteoforms by the single gene. In *K. marxianus BAT1* gene is transcribed into two different mRNAs from two Transcription Start Sites (TSS). The long mRNA (BAT1-L) contains a N-terminal extension (NTE) sequence that is a putative mitochondrial targeting signal (MTS), so BAT1-L encodes a mitochondrial protein. The short mRNA (BAT1-S) encodes an identical polypeptide that lacks the MTS, producing the cytosolic proteoform. The translation initiation codons are represented as +1.

### Localisation of Bat1p depends on the presence of the mitochondrial targeting signal

To understand whether the presence of the MTS determines the localisation of Bat1, fusions of the N-terminal region of Bat1-L and Bat1-S with *GFP* were made and the localisation tested when cells were grown in minimal medium with either ammonium or BCAA (a mixture of valine, isoleucine and leucine (VIL)) as the nitrogen source. For this experiment, expression was from a constitutive promoter, therefore any effects should be independent of growth medium. Cells were treated with the fluorescent probe MitoTracker red and then examined by fluorescence microscopy to visualise of mitochondria and the Bat-GFP fusions (Figure 3). In the figure, mitochondria stain red, Bat1-GFP fusions stain green, and overlap is seen as an orange/yellow colour in the merged images. It was found that Bat1-L-GFP localised to the mitochondria in both nitrogen conditions whereas only cytosolic localisation was seen for Bat1-S-GFP. These data demonstrate that the Bat1 MTS is able to localise a protein to the mitochondrion. We next evaluated whether transcripts from the native promoter of *BAT1* (pNatProm-BAT1) would encode proteins that localised to one or both cellular compartments in a nitrogen-dependent way. For this, we made a construct that included an extended *BAT1* promoter fused to *GFP* such that transcription was possible from either of the two TSS that we had identified and, depending on which TSS was used, an in-frame BAT1-GFP fusion with, or without, the MTS would be made. Localisation was determined by fluorescence microscopy under three different nutrient conditions (Figure 4). First, looking at growth with NH_4_ as the nitrogen source it is seen that BAT1-GFP localises to both the cytosol and the mitochondria. This result was unexpected since it had been predicted that under these biosynthetic conditions, only mitochondrial localisation would be seen. In contrast, and in line with predictions when growing on BCAA, localisation was exclusively to the cytosol. A similar result was seen when cells were growing in the rich medium, YPD, with no mitochondrial localisation. Correlating these data to the potential use of aTSS, it suggests that when amino acids are available as the nitrogen source (BCAA/YPD), only the shorter form of BAT1 is made (using the second TSS, whereas when there is a requirement to synthesise amino acids (NH_4_), both TSS are used and the cell makes the long (mitochondrial) and short (cytosolic) forms of Bat1.

**Figure 3.**
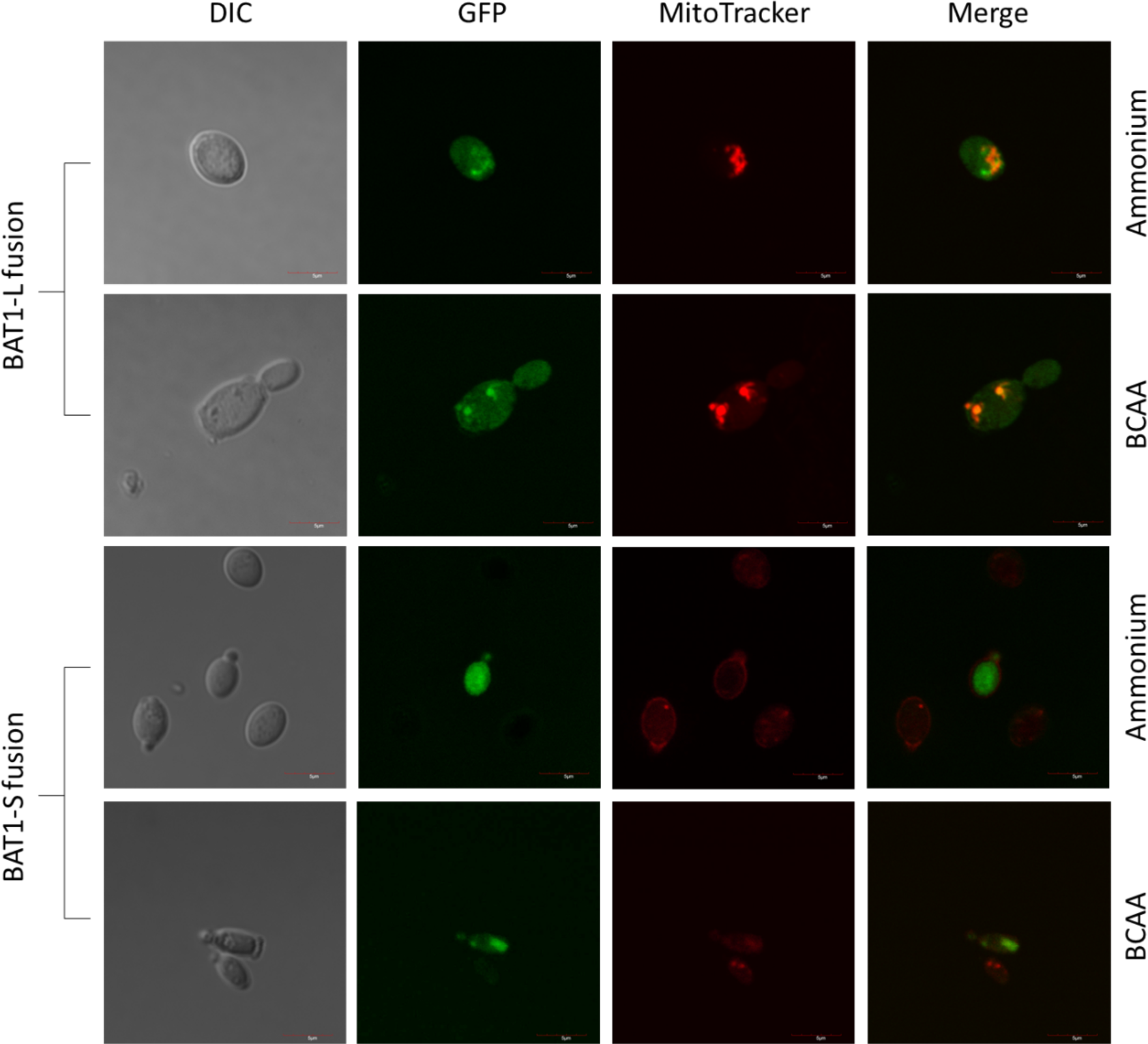
The mitochondrial targeting sequence (MTS) of Bat1 is responsible for mitochondrial localization. Cells transformed with BAT1-L and BAT1-S GFP fusions expressed from the constitutive *TEF1* promoter were grown on ammonium or BCAA, stained with MitoTracker Red and imaged with fluorescence microscopy. The first two rows correspond to cells transformed with BAT1-L fusion carrying the MTS and the second two rows correspond to cells transformed with BAT1-S. From left to right are the images from Differential Interference Contrast (DIC) light, Green Fluorescence Protein (GFP) fluorescence, MitoTracker fluorescence and the merge of GFP and MitoTracker fluorescence. Orange/yellow fluorescence merged images is seen in cases of co-localisation.

**Figure 4.**
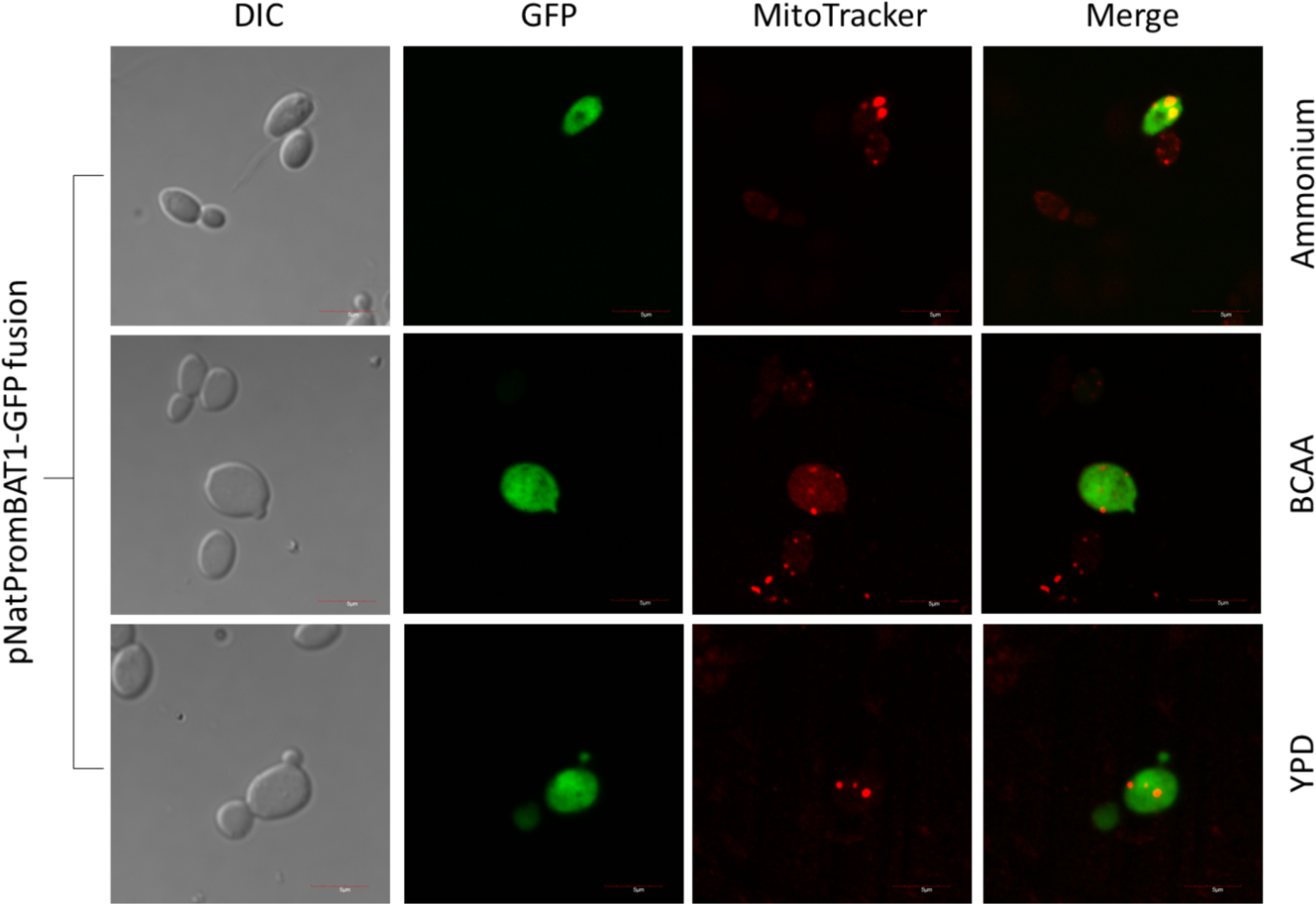
The *BAT1* promoter determines localisation of NTE-GFP in *K. marxianus*. Cells transformed with the pNatProm-BAT1-GFP fusion were grown on ammonium, BCAA or YPD, stained with MitoTracker Red and imaged with fluorescence microscopy as described in figure 3. From left to right are the images from Differential Interference Contrast (DIC) light, Green Fluorescence Protein (GFP) fluorescence, MitoTracker fluorescence and the merge of GFP and MitoTracker fluorescence. Orange/yellow fluorescence is seen when the NTE-GFP protein localises in the mitochondria (Ammonium).

### Expression of different *BAT1* isoforms depending on the nitrogen conditions

To explore the proposed use of aTSS under different nitrogen conditions, the wild-type strain was grown in ammonium and VIL separately and BAT1 mRNA transcripts quantified by RTqPCR. One pair of primers within the *BAT1* open reading frame allowed quantification of the total amount of BAT1 mRNA – BAT1-Total. A second pair specifically amplifies the longer mRNA isoform – BAT1-L, and the levels of BAT-S deduced from the difference between the total amount of BAT1 mRNA and the level of BAT-L. Constitutively expressed *ACT1* was used as a reference and the relative levels of the different isoforms determined when the cells were growing on either ammonium or VIL as a nitrogen source (Figure 5). The overall amount of BAT1 mRNA present in cells growing on VIL was approximately 60% of that seen on ammonium but the major differences arose when comparing the relative amounts of the two isoforms. When growing on ammonium, there were approximately equal amounts of both isoforms whereas on VIL, the shorter isoform (BAT1-S) was almost exclusively present. These data indicate that when growing on BCAA, only the second TSS is used, whereas on ammonium, both TSS seem to be used in approximately equal measure.

**Figure 5.**
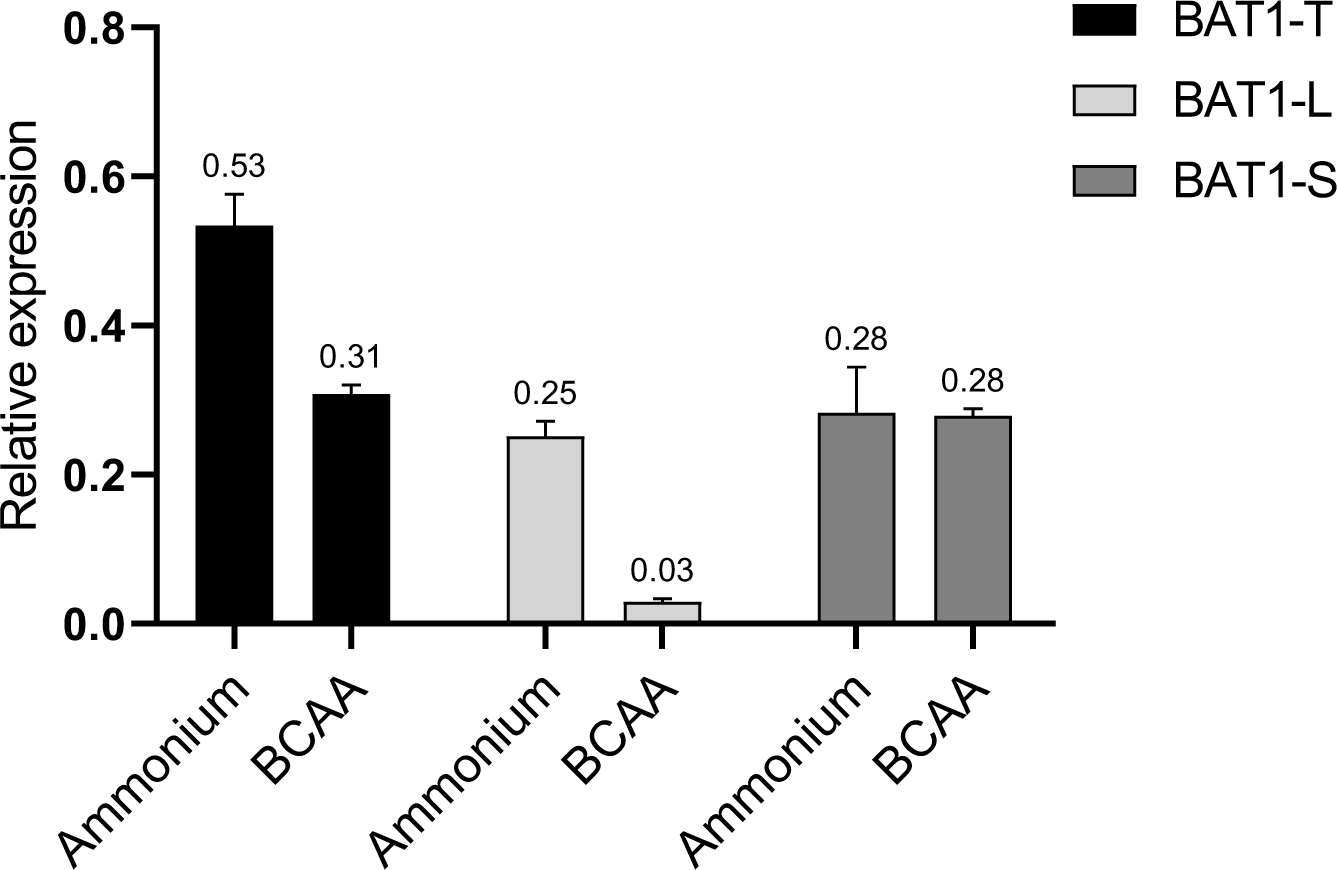
Expression the *BAT1* long isoform (BAT1-L) is decreased when cells are grown on BCAA. Levels of BAT1-L and total BAT1 (BAT1-T; that includes both the long and short isoforms) were measured by RT-qPCR from *K. marxianus* grown on ammonium and BCAA (valine, isoleucine and leucine) and used to also determine the level of BAT-S. The gene expression values are relative to the expression of *ACT1*. Error bars were calculated from biological triplicates.

### Role of the transcription factors in the expression of the long isoform

We next sought to identify the transcriptional factors (TFs) responsible for the use of aTSS in *BAT1*. Two approaches were taken to generate a list of candidate TFs. First, we included TFs that had been shown to be involved in regulation of either *BAT1* or *BAT2* in *S. cerevisiae* (González et al., 2017). Second, we examined the *BAT1* promoter using the *S. cerevisiae* consensus and found potential binding sites that fully matched the consensus for Gcn4, Gln3/Gat1, Hap2, Mot3 and Nrg1, and that deviated by only one nucleotide for Leu3 and Put3. Using a CRISPR-Cas9 method, we created eight deletion mutants, each lacking one of these TFs. The mutants were grown in ammonium and VIL separately and the levels of total BAT1 mRNA and BAT1-L were determined to establish which, if any, of these TFs played a role in expression (Figure 6). For this analysis, the BAT1 mRNA levels in each mutant was compared to that of the wild-type growing under the same conditions and there were clear indications of the involvement of multiple TFs. Looking first at ammonium, it is seen that deletion of *GCN4* had the greatest effect. The total amount of BAT1 mRNA was reduced by about 50%, with essentially none of the longer form made (BAT1-L). Given the earlier data that there is approximately equal transcription from both the first and the second TSS under these conditions (Figure 5), this result indicates that Gcn4 is required for transcription from the first TSS but not the second. Deletion of *LEU3* led to a 50% reduction in BAT-L, showing that Leu3 activates expression from the first TSS but it is not absolutely required. On BCAA, there were visible increases in the total amount of BAT1 in some mutants, though this did not pass the threshold for statistical significance. Looking specifically at use of the first TSS however, a substantial increase in the amount of the long isoform was seen in three mutants: *put3, nrg1* and *hap2.* Although fold-increases were high, they did not dramatically change the total amount of BAT1 mRNA because the level of BAT1-L was coming from a very low base in the wild-type (Figure 5). Nevertheless, the conclusion is that all three TFs impair expression from the first TSS on BCAA. Since most *BAT1* expression on BCAA is of the short isoform, these data suggest that none of these TFs are required for expression from the second TSS on BCAA.

**Figure 6.**
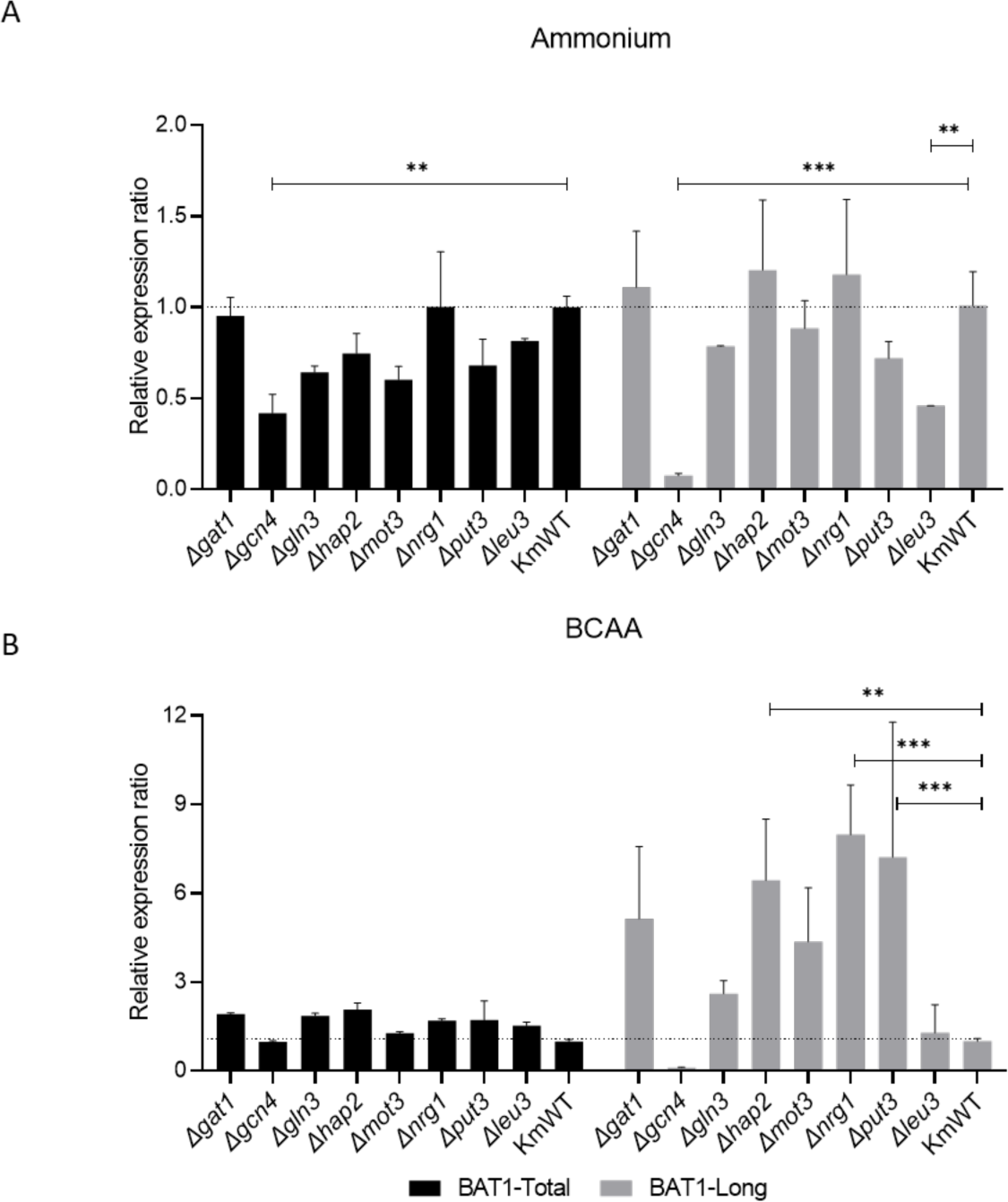
Role of transcription factors in the expression of *BAT1*. Levels of BAT1-L and total BAT1 (BAT1-T; that includes both the long and short isoforms) were measured by RT-qPCR from the indicated *K. marxianus* wild-type (WT) and mutant strains grown on ammonium (panel A) and BCAA (valine, isoleucine and leucine)(panel B). Data are normalised to ACT1 mRNA levels and the ratio of BAT1-L and BAT1-T in each mutant relative to wild type is shown. Data represent two independent experiments with technical triplicates. The differences were considered statistically significant when the p-values were <0.001 (***) or <0.01 (**).

### Function of Bat1p in *K. marxianus* depending on the presence of the NTE

The data are consistent with the hypothesis that when growing on ammonium (biosynthetic conditions), both proteoforms are made, with the longer form localising to the mitochondrion to enable synthesis of BCAA. On VIL as a nitrogen source (catabolic conditions), the mitochondrial proteoform would not be required and cytosolic Bat1 is involved in degradation of the BCAA to provide nitrogen for the cell. To explore this further, we constructed two mutants of *BAT1*, one completely inactivating the gene (*Δbat1*) and one precisely deleting the MTS so that only the cytosolic proteoform is made (*ΔNTE-bat1*). These mutants were then grown on different sole nitrogen sources and the maximum specific growth rates (μ_max_) were calculated (Figure 7). The *Δbat1* mutant showed severely impaired growth on biosynthetic conditions (ammonium and glutamine) and on individual amino acids, but only moderately impaired growth in catabolic conditions (BCAA). When provided with both ammonium and BCAA, growth of *Δbat1* was restored almost to the wild-type rate confirming that Bat1 is required for both the anabolism and catabolism of BCAA. Somewhat unexpectedly, the *ΔNTE- bat1* strain did not display any growth impairment on ammonium, indicating that BCAA biosynthesis is possible even in the absence of the mitochondrially-targeted protein. In fact, only discernible phenotype for *ΔNTE-bat1* was a reduction of about 50% in growth rate on BCAA.

**Figure 7.**
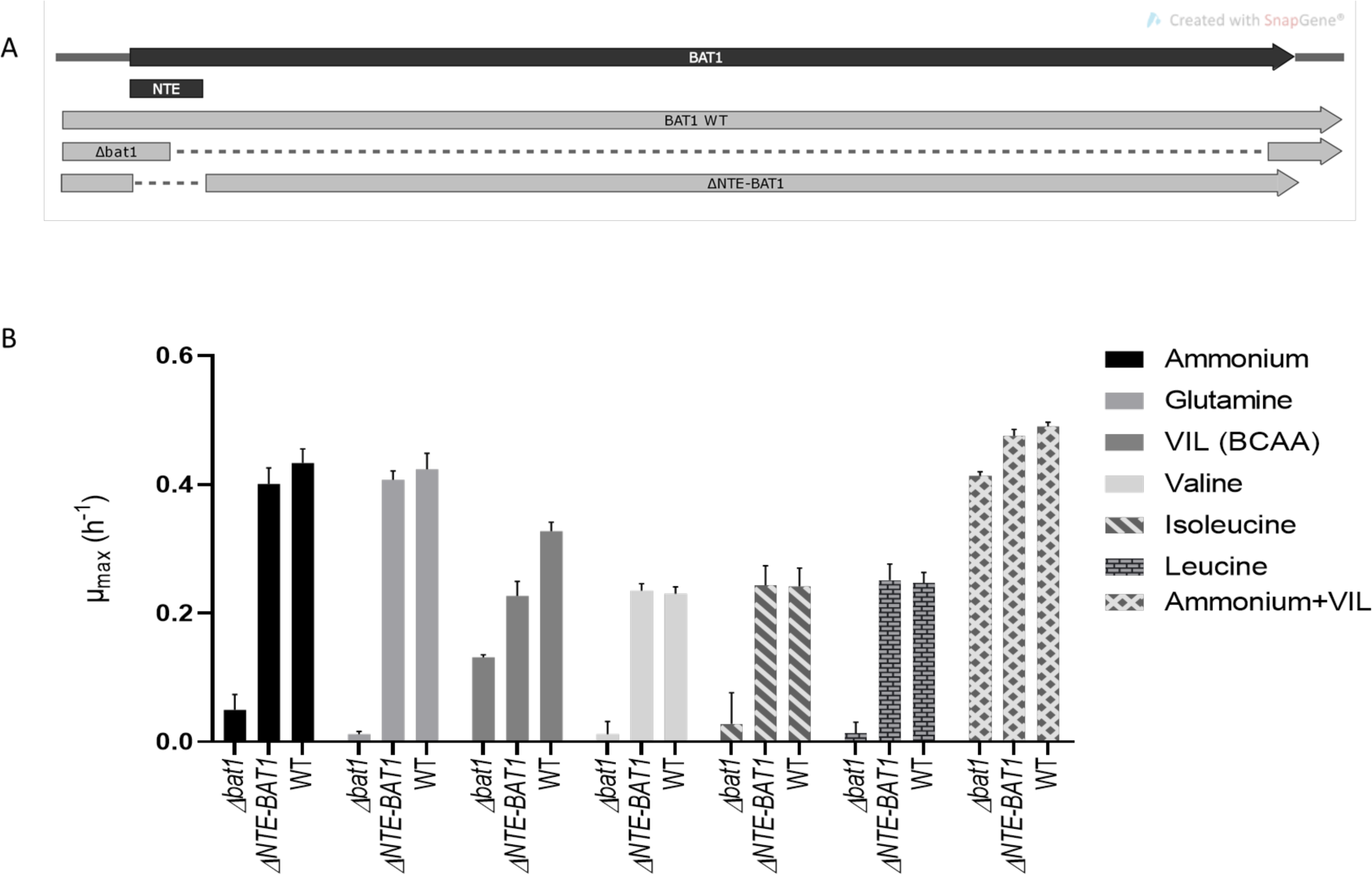
Growth rates of *K. marxianus* bat1 mutant strains. Two mutant strains, one lacking the B*AT1* coding region (D*bat1*) and one with a precise in-frame deletion of the mitochondrial targeting sequence (DNTE-*BAT1*) were generated and growth rates on different nitrogen sources were determined. (A) Schematic depiction of the *K. marxianus* mutants. (B) Bar graph of maximum specific growth rate (μ_max_) of mutants and wild type (WT) grown in the indicated nitrogen sources. Data were compiled from four independent experiments with technical replicates.

### Wider use of aTSS for controlling sub-cellular localisation

The data with *BAT1* raise the question as to whether the mechanism of use of alternative TSSs for the production of two protein isoforms with different sub-cellular localisation might be more widespread. To explore this, we used the GWIPs platform to examine two other *K. marxianus* genes, *ALT1* and *BAP3*, that may also need to encode differentially located proteoforms. In *S. cerevisiae*, *ALT1/ALT2* (Peñalosa-Ruiz et al., 2012) and *BAP2/BAP3* (Regenberg et al., 1999) are paralogous pairs, one of which encodes a mitochondrially-localised proteoform. In both cases, we found that the *K. marxianus* orthologue has more than one TSS and a second in-frame AUG (Figure 8). Under the conditions that the data was generated, there is unambiguous translation from first AUG (based on the ribosome profiling footprints), but the major transcription start site is downstream of this. The most straightforward explanation is that, as with *BAT1*, alternative transcription starts sites are used to generate mRNAs that give rise to proteins with alternative N-termini. For *ALT1*, there is a relatively short N-terminal extension that encodes a canonical MTS (TargetP 2.0 score 0.993), and it is clear that differential use of the aTSS can give rise to the alternative proteoforms that will differentially localise in the exact same manner as with *BAT1*. *BAP3* is somewhat different since the difference in length between the two proposed proteoforms is 71 amino acids and an MTS is not evident in the N-terminal extension. Bap2/Bap3 proteoforms localise to the plasma-membrane, ER and mitochondrion in *S. cerevisiae* (Regenberg et al., 1999), and it is possible that the localisation uses an alternative mechanism (such as protein folding) to the canonical MTS. Although more experimental data would be required to elucidate details, taking *BAT1, ALT1* and *BAP3* as examples, it can be inferred that the use of aTSS to direct the synthesis of differentially-localised proteoforms is common.

**Figure 8.**
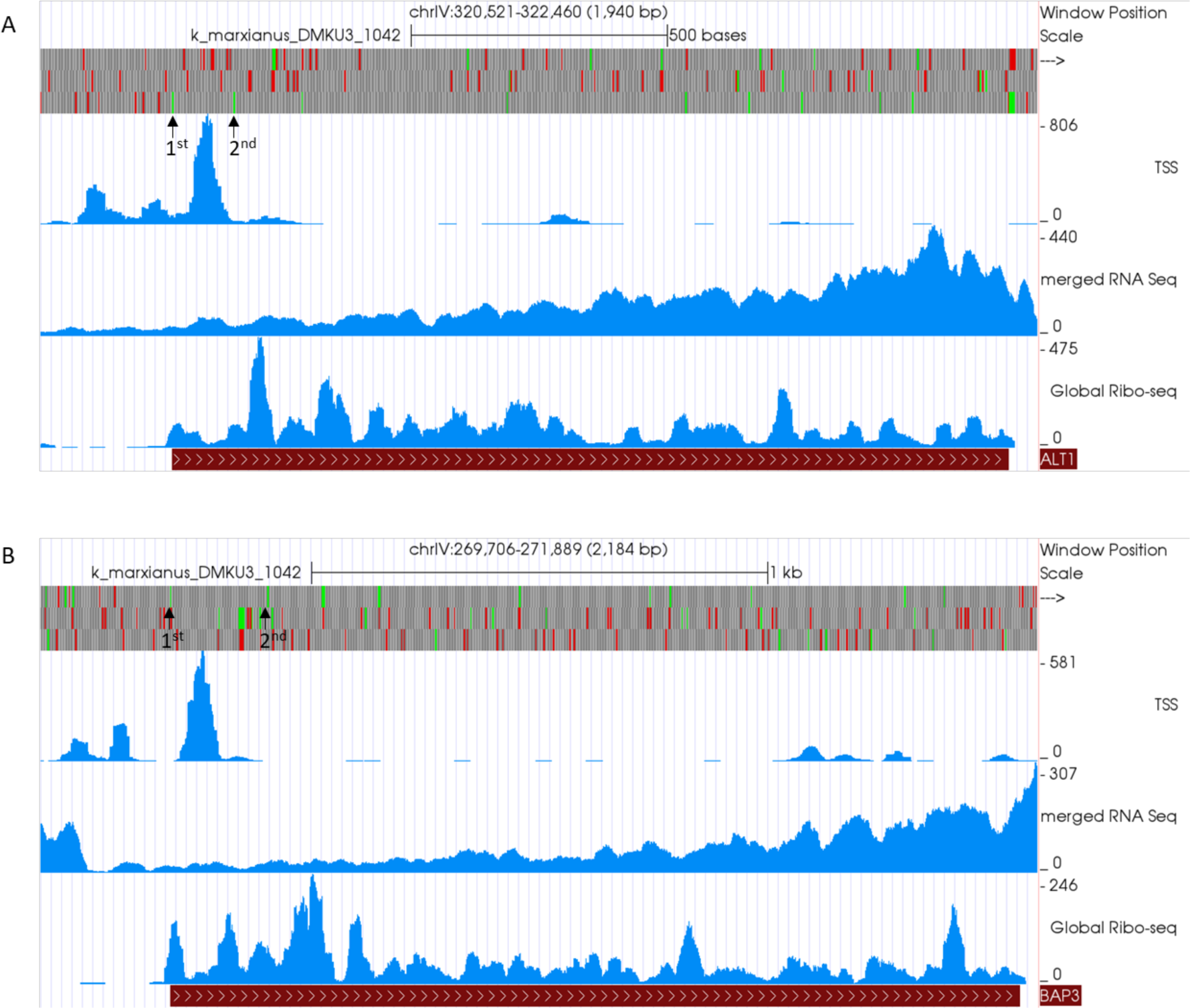
Visualisation of the *ALT1* and *BAP3* loci in the *K. marxianus* GWIPS-Viz Genome Browser. Panel 1, *ALT1*; panel B, *BAP3.* The red bar at the bottom represents the coding region of each gene and the top tracks depicts the ORF architecture including start codons in green and stop codons in red (*ALT1*, 3^rd^ track; *BAP3*, 1^st^ track). Note that scales are not identical. The two black arrows indicate the positions of the first and the second in-frame start codons of each gene Coverage of Ribosome profiling (Ribo-Seq), RNA Sequencing (RNA- Seq) and Transcription Start Site Sequencing (TSS-Seq).

## DISCUSSION

The increased availability of omics datasets and on-line databases for non-traditional or non-conventional yeasts are valuable assets for the investigation of the evolution of gene regulatory mechanism. We exploited these resources to perform an *in silico* analysis of the gene encoding branched chain aminotransferases (BCAT) in *K. marxianus*, a species that diverged from the *Saccharomyce*s lineage prior to the whole genome duplication (WGD). Previous studies in the related species *K. lactis* and *L. kluyveri* had established that *BAT1* is bifunctional, encoding both the mitochondrial and cytosolic BCAT, required for synthesis or degradation of branched chain amino acids (BCAA), respectively. It was proposed that synthesis of the required proteoform was dependent on the nitrogen source in the growth medium, but the underlying mechanism was not determined. In principle, there are a number of different means that yeast could use to generate these alternative proteoforms. For example, a global analysis of the transcriptional and translational landscape in *K. marxianus* found that proteins with alternative N-termini could be generated through the use of alternative transcriptional start sites (aTSS) or leaking scanning of a single transcript, whereby an AUG in a poor context (or a non-cognate start codon) is bypassed enabling initiation at a downstream AUG codon (Fenton et al., 2022b). Another system is seen with the *S. cerevisiae* fumarase Fum1, which uses alternative protein folding to control the distribution of this enzyme between the mitochondrion and cytosol (Herrmann, 2009; Regev-Rudzki et al., 2009). By analysing transcription start sites (TSS), transcriptome data, and ribosome profiling data, we formed a hypothesis that *K. marxianus* uses aTSS to produce cytosolic or mitochondrial proteoforms of BCAT. We proposed that when biosynthesis of BCAA was required, transcription would initiate at the first TSS giving rise to a longer proteoform containing a mitochondrial targeting signal (MTS), whereas when BCAA are in excess or providing the cell with a source of nitrogen, the second TSS would be used, producing a shorter cytosolic proteoform. Using GFP fusions, we confirmed both the presence of an MTS and nitrogen-source differential localisation of Bat1. Finally, we directly measured levels of mRNA isoforms that would arise from use of the first or the second TSS and found that when the yeast is growing using BCAA as a N-source, there is essentially no transcription from the first TSS – only the second TSS is used and therefore only the cytosolic proteoform is made. In contrast, under biosynthetic conditions (NH_4_ as a N-source), both transcription start sites are used in almost equal measure leading to synthesis of mitochondrial and cytosolic proteoforms.

Having established that synthesis of the proteoforms is controlled at a transcriptional level, we used a combination of *in silico* analysis of the *BAT1* promoter and knowledge from *S. cerevisiae* to identify transcription factors (TFs) that could be responsible for the transcriptional control. Eight candidate TFs were individually inactivated by CRISPR-Cas9 – mediated deletion and the levels of the mRNA isoforms measured under different conditions. The strongest effect was seen for Gcn4, which appears to be absolutely required for transcription from the first TSS when the cells are growing under biosynthetic conditions (NH_4_). The *K. marxianus BAT1* promoter harbours the consensus binding sequence of Gcn4p ∼59 bp upstream the first TSS, ideally located for direct binding of Gcn4 to drive expression. This accords with data from *S. cerevisiae* where it has long been known that Gcn4 is a transcriptional activator of amino acid biosynthetic genes, responding to amino acid starvation by a translational control mechanism (Hinnebusch, 2005; Natarajan et al., 2001). Furthermore, in *S. cerevisiae*, Gcn4 directly binds to the *BAT1* promoter and activates expression only in biosynthetic conditions (Gonzalez et al., 2017), a situation that is analogous to transcription of the longer mRNA isoform in *K. marxianus*. Although, less pronounced, Leu3 is also required for full activation of expression from the first TSS when cells are growing under biosynthetic conditions (NH_4_). This also mirrors *S. cerevisiae* where Leu3 activates expression of *BAT1*, though in this case, Leu3 is considered the major activator. The candidate Leu3 binding sites in the *K. marxianus BAT1* promoter deviate by a single nucleotide from the consensus, and are quite distal from the transcription start site. One or both of these factors may explain the apparent difference in importance of Leu3 in the two yeasts.

Looking at catabolic conditions (BCAA), we found roles for three TFs. Interestingly, none of these was a positive effect since the data indicate that Put3, Nrg1 and Hap2 inhibit transcription from the first TSS on VIL. Nrg1 works by recruiting the Ssn6-Cyc2 co-repression complex and it is possible that this TF represses transcription by recruiting this complex to the *BAT1* promoter (Murad et al., 2001; Park et al., 1999). Hap2 forms part of the global regulator HAP complex, binding the 5’-CCAAT-3’ sequence. This conserved binding site is found in 30% of eukaryotic genes and tends to be located at 60 to 100 bp from the TSS (Dolfini et al., 2009) whereas in the *BAT1* promoter, it is 450 bp upstream from the first TSS. Possibly the HAP complex may interact with Nrg1 to recruit the Ssn6-Cyc2 co-repressors or, alternatively, the effect may be indirectly mediated via another gene, itself regulated by the HAP complex. In *S. cerevisiae,* under catabolic conditions, Put3 binds to the *BAT2* promoter, activating expression and is believed to also repress expression of *LEU1*, reducing the levels of α-iPM and essentially switching the role of Leu3 at the *BAT1* promoter from an activator to a repressor (González et al., 2017). The *K. marxianus BAT1* promoter has a sequence that differs from the Put3 consensus by one nucleotide so in principle Put3 binding to the promoter is possible. Interestingly, the *K. marxianus LEU1* promoter lacks a consensus Put3 binding site so it is possible that the negative effect of Put3 on transcription from the first TSS is a direct effect. In contrast to *S. cerevisiae BAT2*, there is no evidence that Put3 activates expression from the second TSS in *K. marxianus*. There is also no evidence for a role for Gln3. In *S. cerevisiae*, expression of *BAT2* involves direct binding of Gln3, the classic TF involved in the NCR system. This is not the case in *K. marxianus* since there is not a loss of transcription from the second TSS in the *gln3* mutant. Although a putative Gln3 binding site is present in the *Kluyveromyces BAT1* promoter, it is far from the transcription start sites and unlikely to be used. In fact, this is similar to the *S. cerevisiae BAT1* promoter where deletion of putative Gln3 binding sites had no effect (González et al., 2017).

One of the goals of this study was to address the hypothesis that retention of both BCAT paralogues in *S. cerevisiae* offers greater regulatory control than in the ancestral state where both proteoforms are synthesised from a single gene. Our data support that hypothesis, since we showed that while the pre-WGD form of the gene is capable of differential synthesis of the two proteoforms, the regulatory control appears to be less tight than that of the post-WGD form. This conclusion is based on multiple data showing that under biosynthetic conditions both proteoforms are made in *K. marxianus*, while, in *S. cerevisiae*, only the mitochondrial proteoform is made. Under catabolic conditions, equal resolution is achieved. Mechanistically, the difference can be explained by the fact that it is easier to control the expression of two distinct genes than the use of two transcription start sites 75 nucleotides apart. Based on our data, we propose that the ancestral gene largely controlled expression ratios via regulation at the first TSS – either activating via Gcn4 and Leu3 or repressing via Put3 or Ssn6-Tup1 (recruited by Nrg1) (Figure 9). Most of these mechanisms were retained in *BAT1* following the WGD, though the Put3 regulon expanded to include *LEU1* offering an additional means to repress *BAT1*. The pre-WGD expression of the second TSS, giving rise to the cytosolic proteoform, was essentially unregulated but in post-WGD species, *BAT2* lost the MTS and acquired new regulation, specifically by integration into the NCR system (via Gln3) and co-opting Put3 as a second activator by optimising the sub-optimal binding sites. At the same time, the constitutive expression of this proteoform seen in *K. marxianus* was lost, therefore *BAT2* is actually dependent on these activators for expression. One interesting feature of *S. cerevisiae BAT2*, not seen in *BAT1*, is the presence of positioned nucleosomes that occlude some potential binding sites. The gene promoters in both pre- and post-WGD species have binding sites for several TFs associated with chromatin remodelling, which is suggesting of a role for chromatin structure in regulating expression. Although major effects were not seen with relevant single mutants in either *S. cerevisiae* or *K. marxianus*, it is possible that there is redundancy and this would be an interesting area to explore further. Overall, our data supports a process whereby post-WGD yeasts maintained most of the ancestral regulation in one copy of a duplicated gene and adapted existing cis and trans acting factors to develop a new regulatory framework for the second copy.

**Figure 9.**
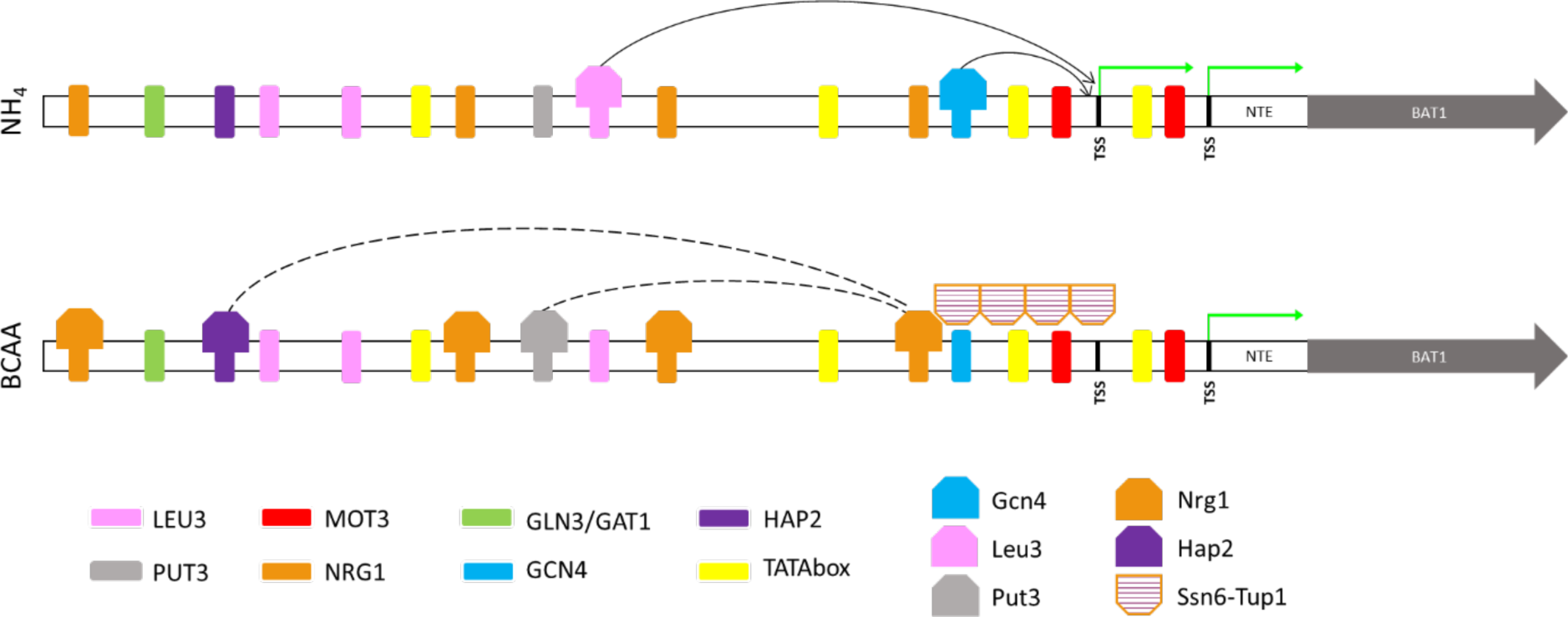
Cartoon with the proposed transcriptional regulation mechanism of *BAT1* in *K. marxianus* when growing under biosynthetic (NH_4_) or catabolic (BCAA) conditions. Potential binding sites in the BAT promoter (∼600 bases upstream of the first AUG are included) for different transcription factors (TF) are show in various colours. Transcription from the first or the second TSS is indicated with a green arrow. When growing on NH_4_, it is proposed that Leu3 and Gcn4 and activate expression from the first TSS. Expression from the second TSS is also possible and so both the cytosolic and mitochondrial forms of Bat1 are made. Only binding to one putative Leu3 motif is shown for simplicity. When growing on BCAA, it is proposed that Nrg1 recruits the Ssn6-Tup1 co-repressor complex, leading to occlusion of the first TSS but allowing use of the second TSS – thereby making the cytosolic form of Bat1. It is suggested that Hap2 and Put3 may contribute to the recruitment of the co-repression complex but that is quite tentative.

## Acknowledgements

We would like to acknowledge the BioSciences Imaging Centre (Department of Anatomy & Neuroscience, BioSciences Institute, University College Cork) for assistance imaging specimens for this research. We also thank Martine Pradal and Thérèse Marlin (INRAE, UMR Sciences pour l’oenologie, Montpellier, France) for helpful technical assistance.

## Funding

This project was funded by the European Union Horizon 2020 research and innovation programme under the Marie Skłodowska-Curie Actions Grant Agreement No. 764927 and Research and Innovation Grant Agreement No. 720824

## Conflict of Interest

P.V.B is a co-founder of RiboMaps Ltd, a company that provides ribosome profiling as a service.

## Supplementary Figures and data

**Supplementary figure 1.**
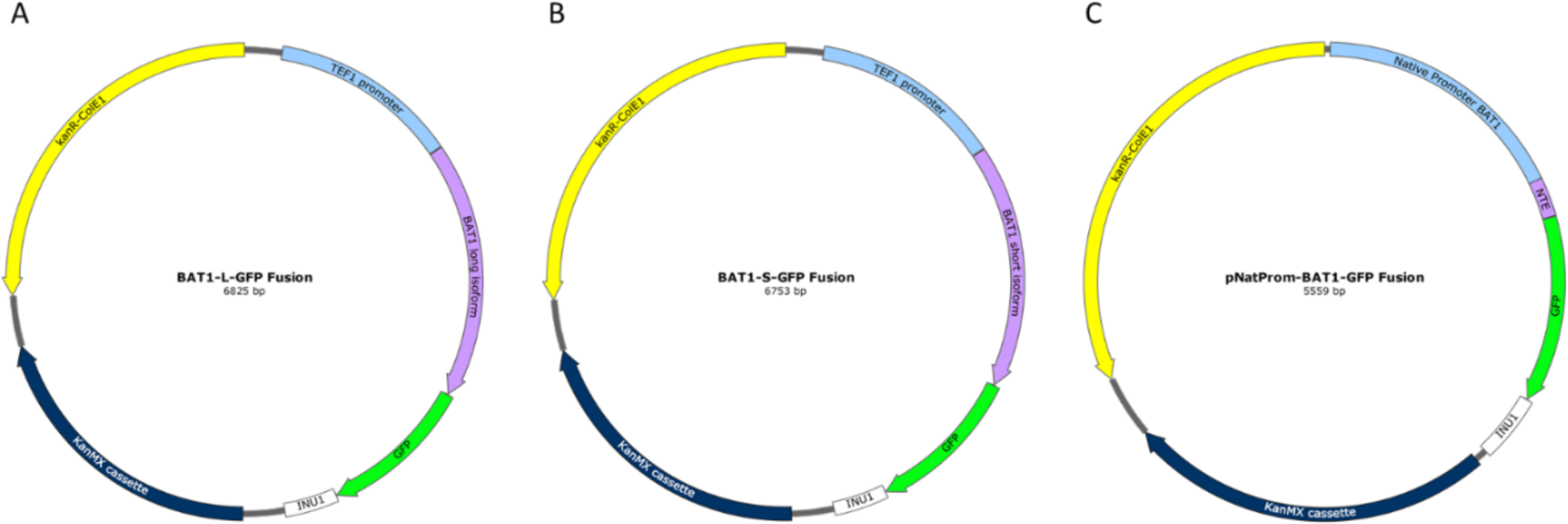
Plasmids constructed for localisation of *BAT1* in *K. marxianus*. (A) BAT1-L fusion constructed for localisation of the long isoform of *BAT1* fused with GFP and induced by *TEF1* promoter. (B) BAT1-S fusion constructed for localisation of the short isoform of *BAT1* fused with GFP and induced by *TEF1* promoter. (C) pNatProm-BAT1 fusion containing the NTE fused to GFP and induced by the endogenous promoter of *BAT1.* All plasmids have *INU1* terminator. KanMX cassette for G418 resistance used for yeast selection and KanR-ColE1 contains the replication origin and selection marker of bacteria with kanamycin.

**Supplementary table 1.**
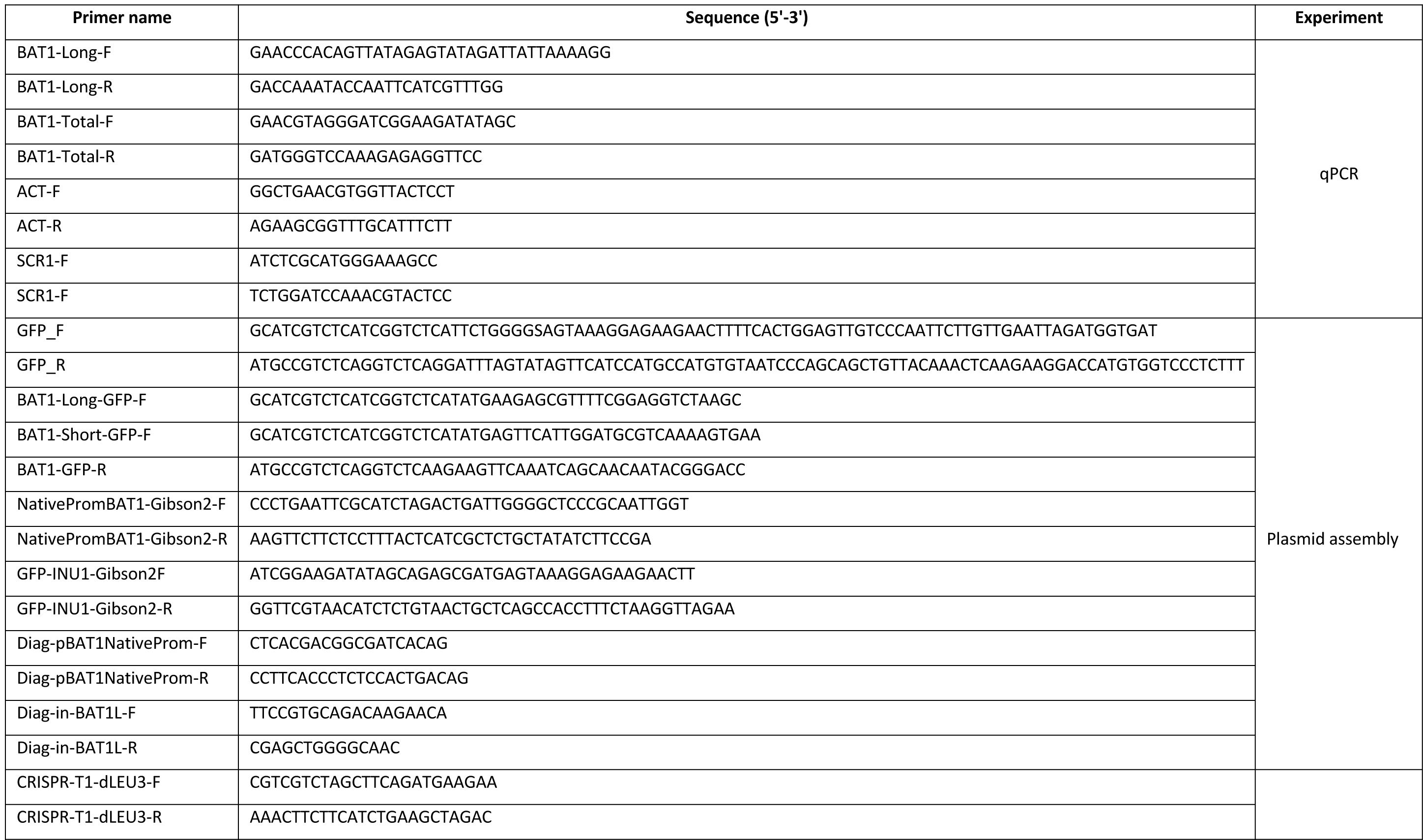
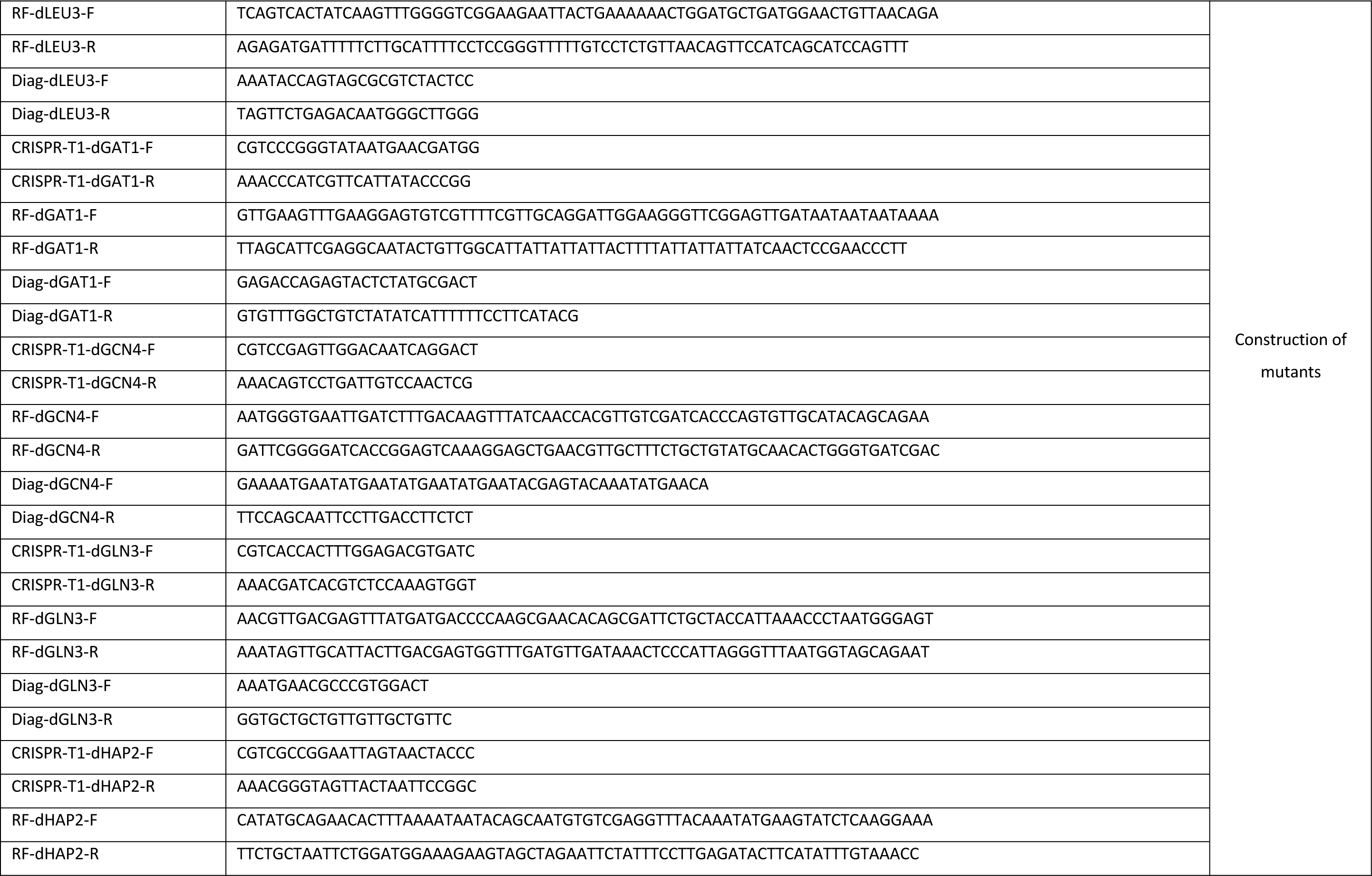
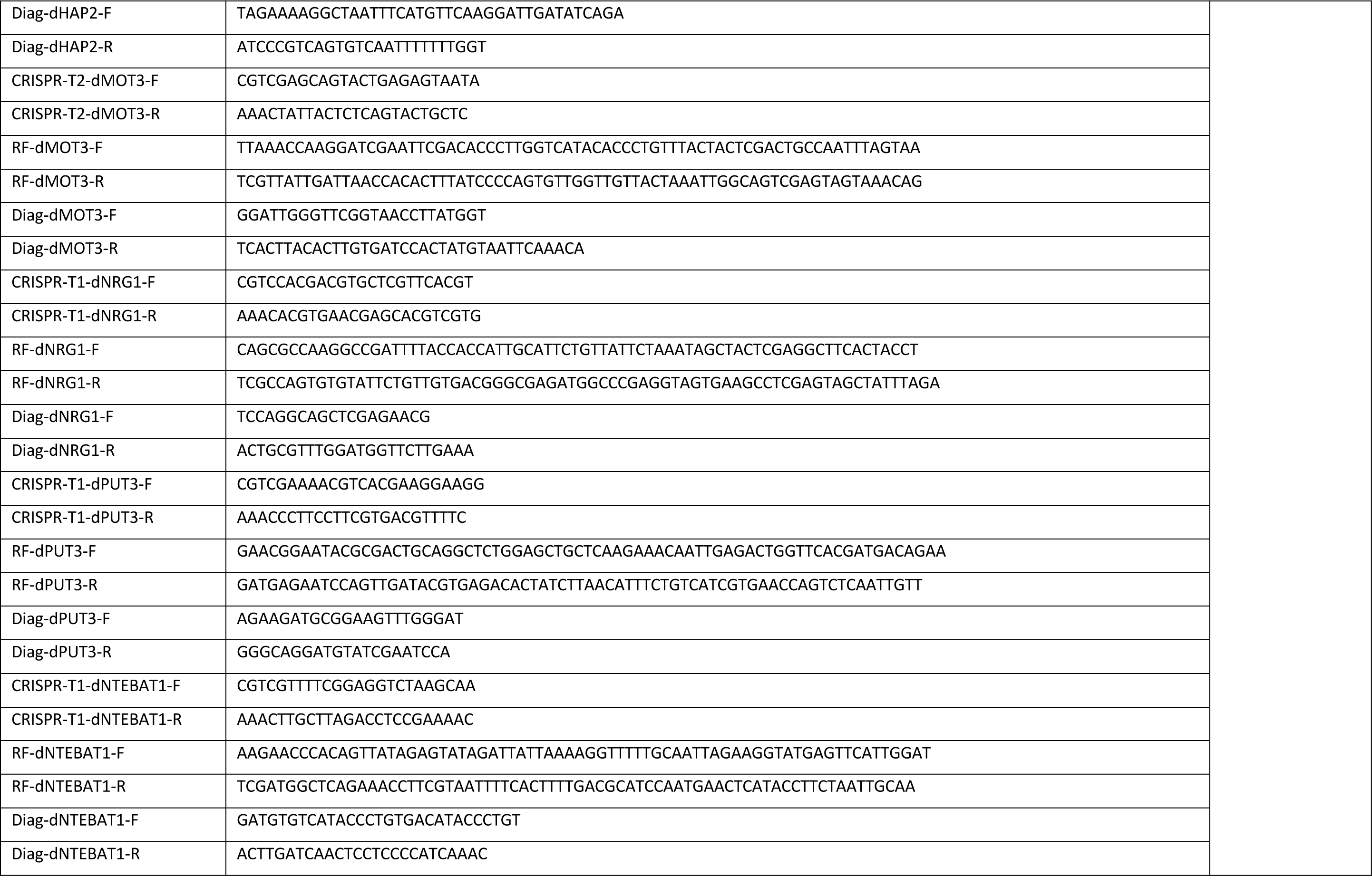
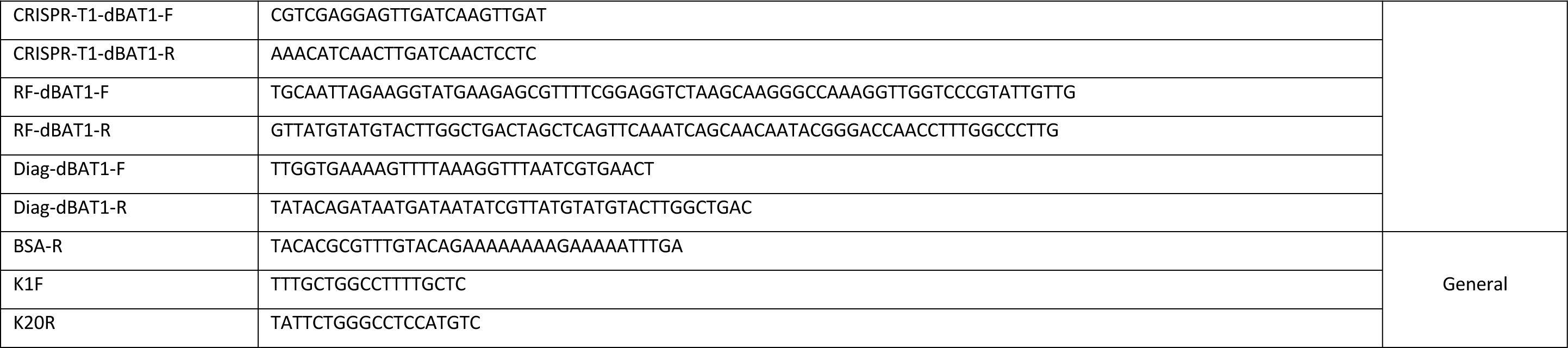
Primers used in this study

Supplementary material describing the construction of the *K. marxianus* transcription factor mutants

Mutants genotype and phenotype:

| Strain name | Genotype of mutants | Deleted sequence (bp) | Growth phenotype |  |  |  |
| --- | --- | --- | --- | --- | --- | --- |
|  |  |  | Ammonium | BCAA | Proline | Galactose |
| *K. marxianus* NBRC1777 $\Delta gat1$ | CHR VI $\Delta 635,534 - 636,959$ | 1426 | Good | Good | - | - |
| *K. marxianus* NBRC1777 $\Delta gcn4$ | CHR V $\Delta 868,076 - 868,697$ | 622 | Poor | Poor | - | Poor |
| *K. marxianus* NBRC1777 $\Delta gln3$ | CHR III $\Delta 1,456,475 - 1,457,877$ | 1403 | Poor | Poor | - | - |
| *K. marxianus* NBRC1777 $\Delta hap2$ | CHR III $\Delta 1,550,967 - 1,551,414$ | 447 | Good | Good | - | - |
| *K. marxianus* NBRC1777 $\Delta mot3$ | CHR VI $\Delta 700,804 - 701,788$ | 984 | Good | Good | - | - |
| *K. marxianus* NBRC1777 $\Delta nrg1$ | CHR IV $\Delta 283,460 - 284,113$ | 653 | Good | Good | - | - |
| *K. marxianus* NBRC1777 $\Delta put3$ | CHR VI $\Delta 343,875 - 345,730$ | 1854 | Good | Good | Null | - |
| *K. marxianus* NBRC1777 $\Delta leu3$ | CHR VI $\Delta 946,030 - 947,790$ | 1760 | Good | Good | - | - |
| *K. marxianus* NBRC1777 $\Delta bat1$ | CHR I $\Delta 1,068,093 - 1,069,223$ | 1130 | Null | Poor | - | - |
| *K. marxianus* NBRC1777 $\Delta NTE-BAT1$ | CHR I $\Delta 1,068,052 - 1,068,127$ | 75 | Good | Good | - | - |
| *K. marxianus* NBRC1777 WT | - | - | Good | Good | Good | Good |

Growth of mutants in different conditions compared to the wild type:

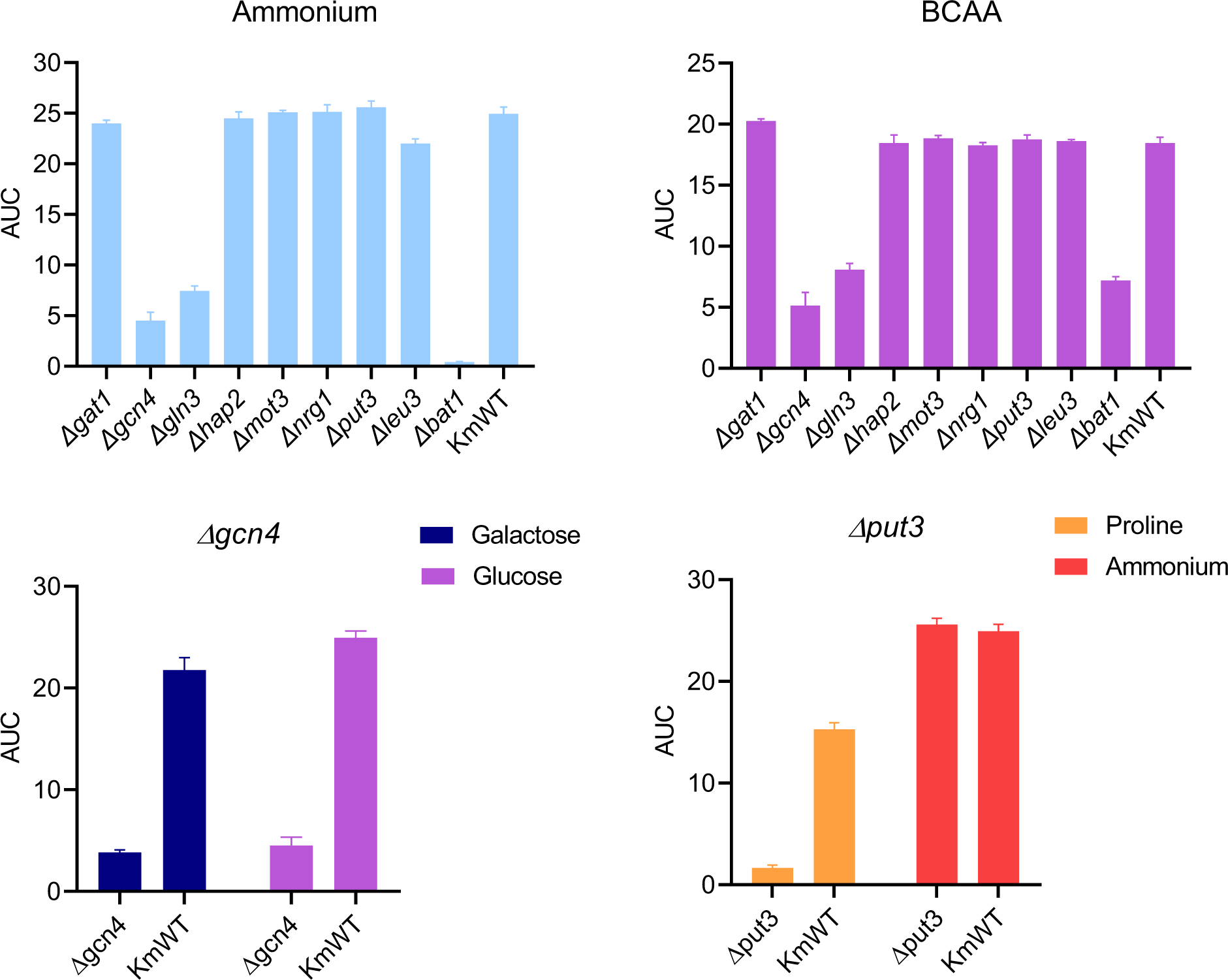

Deletions in the target locus:

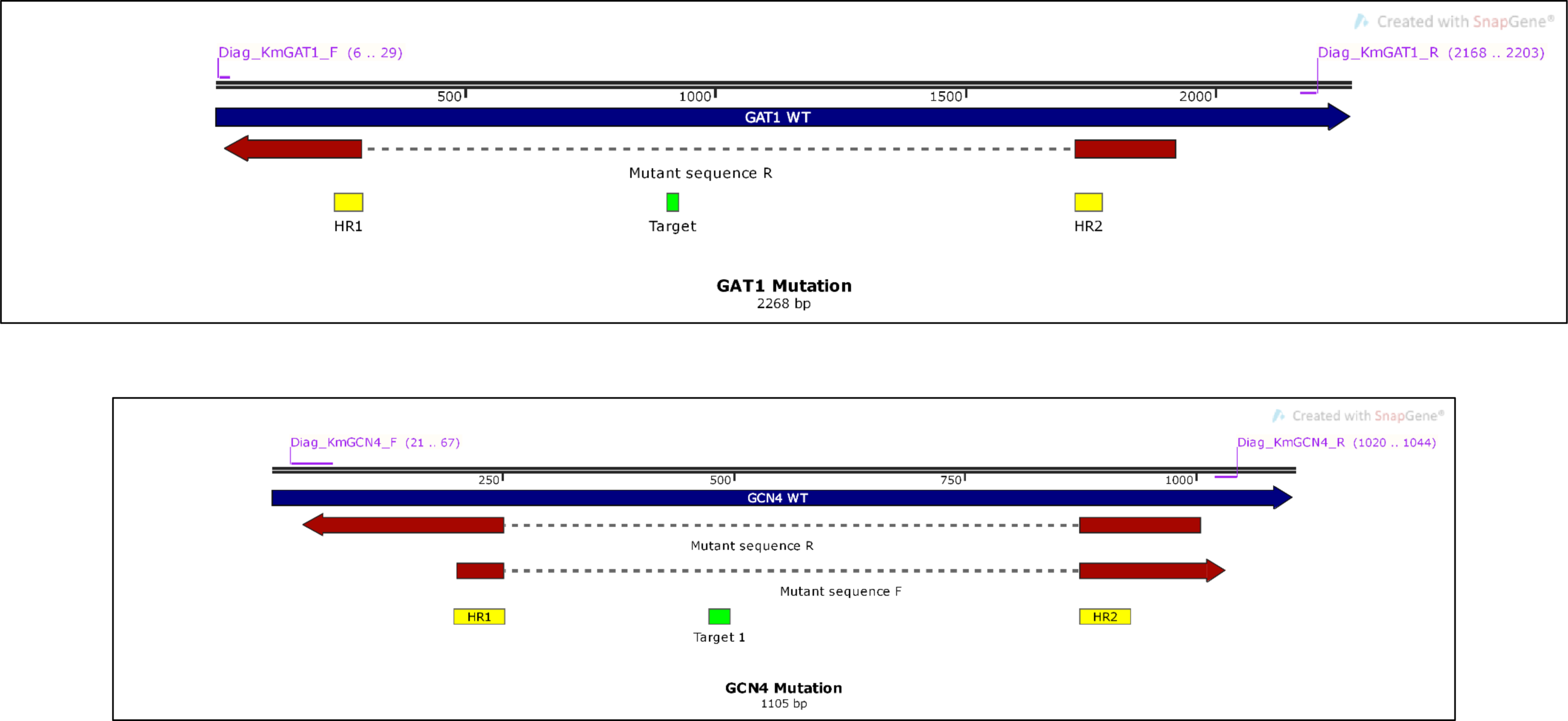
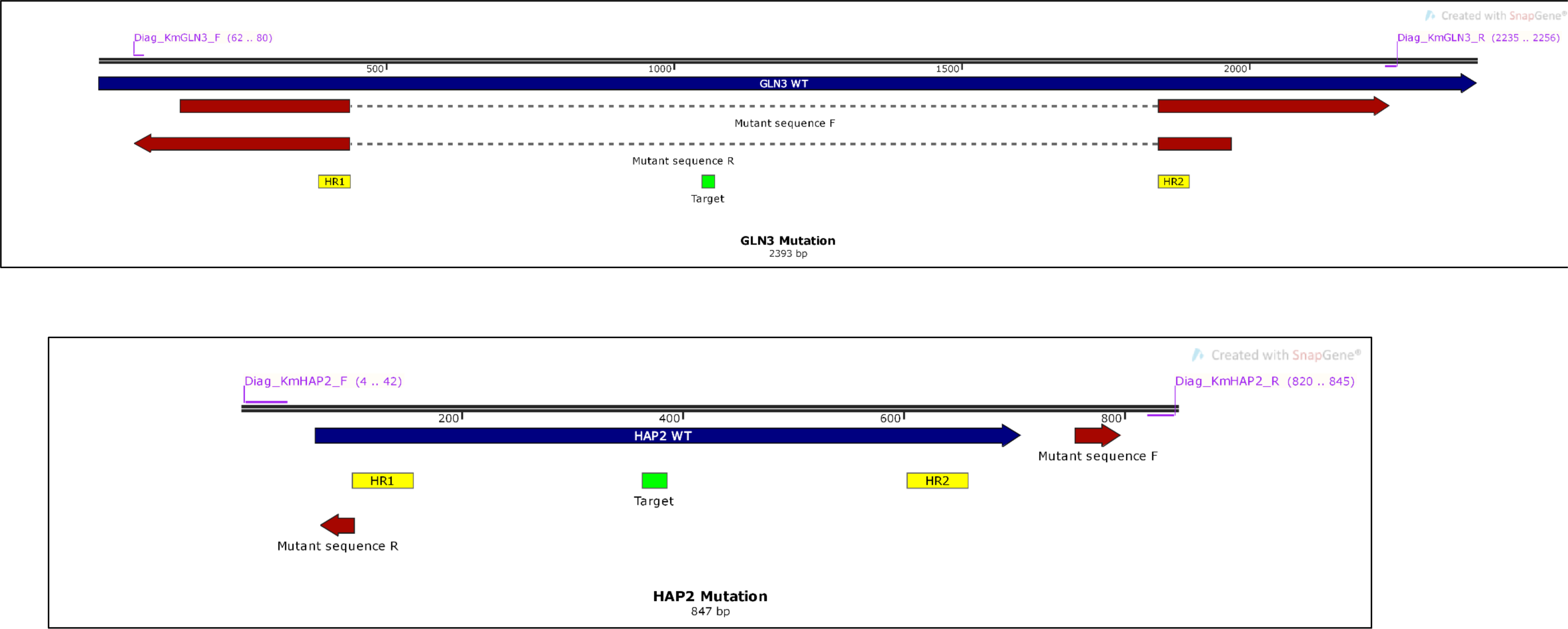
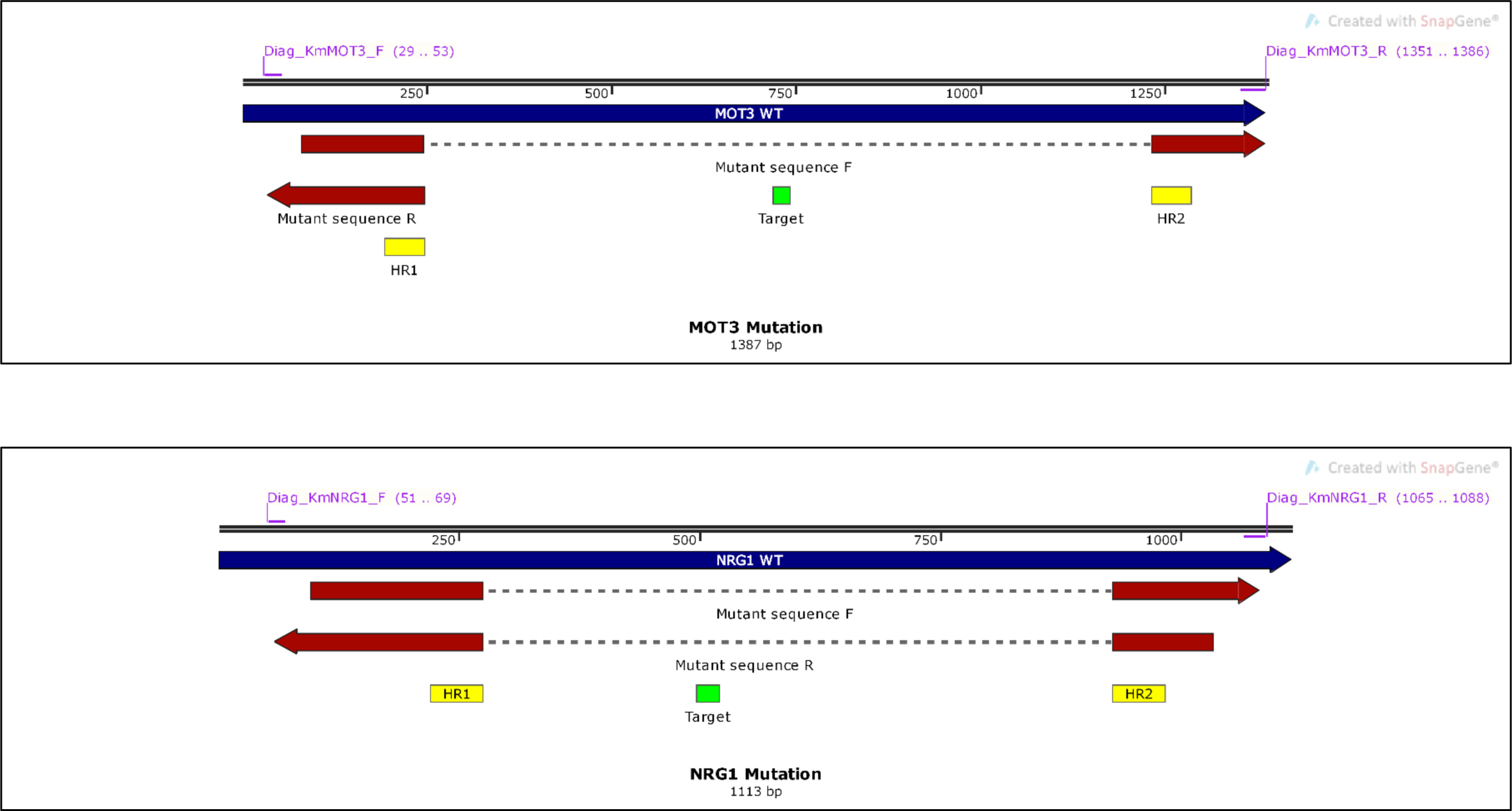
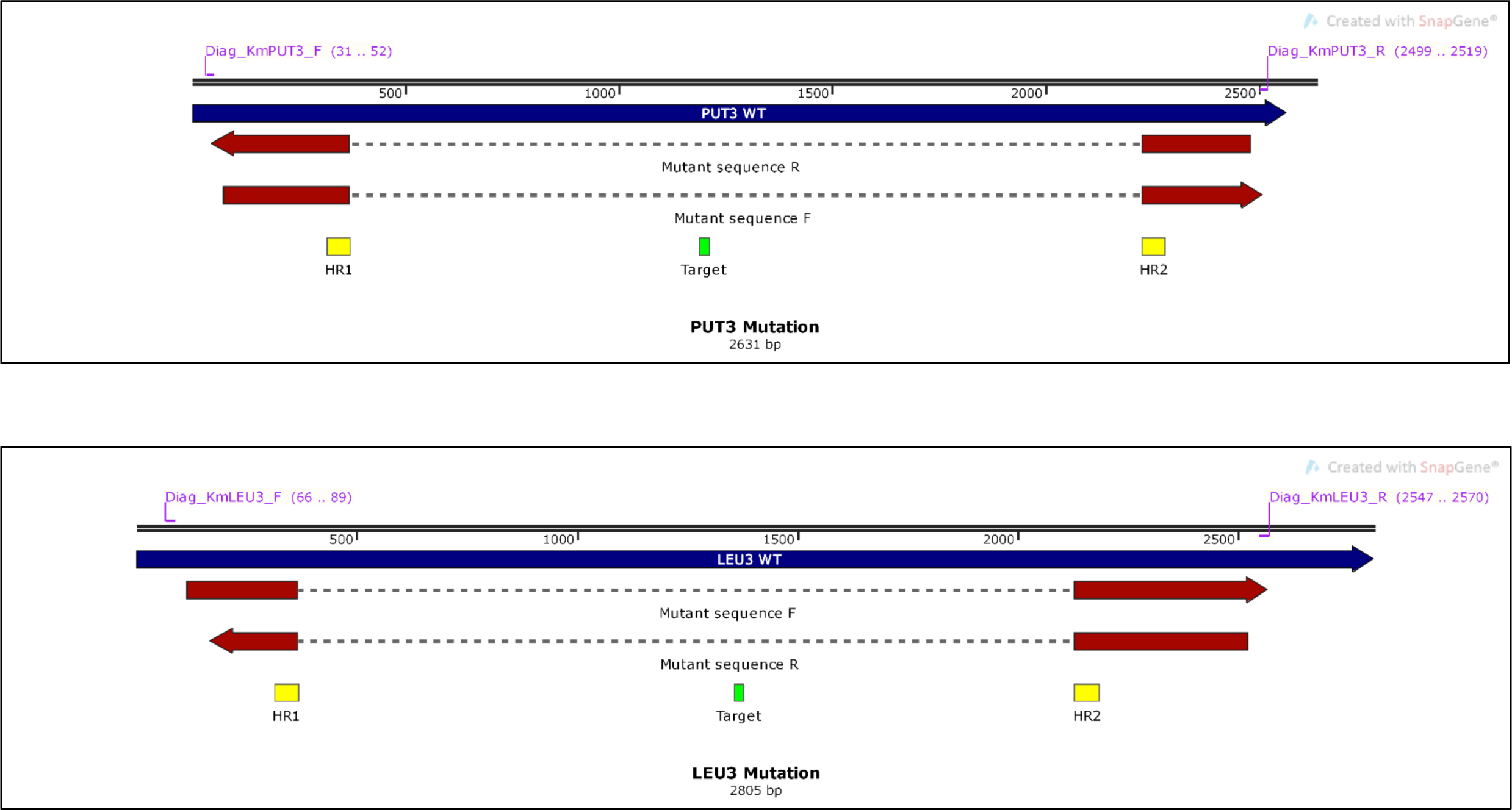
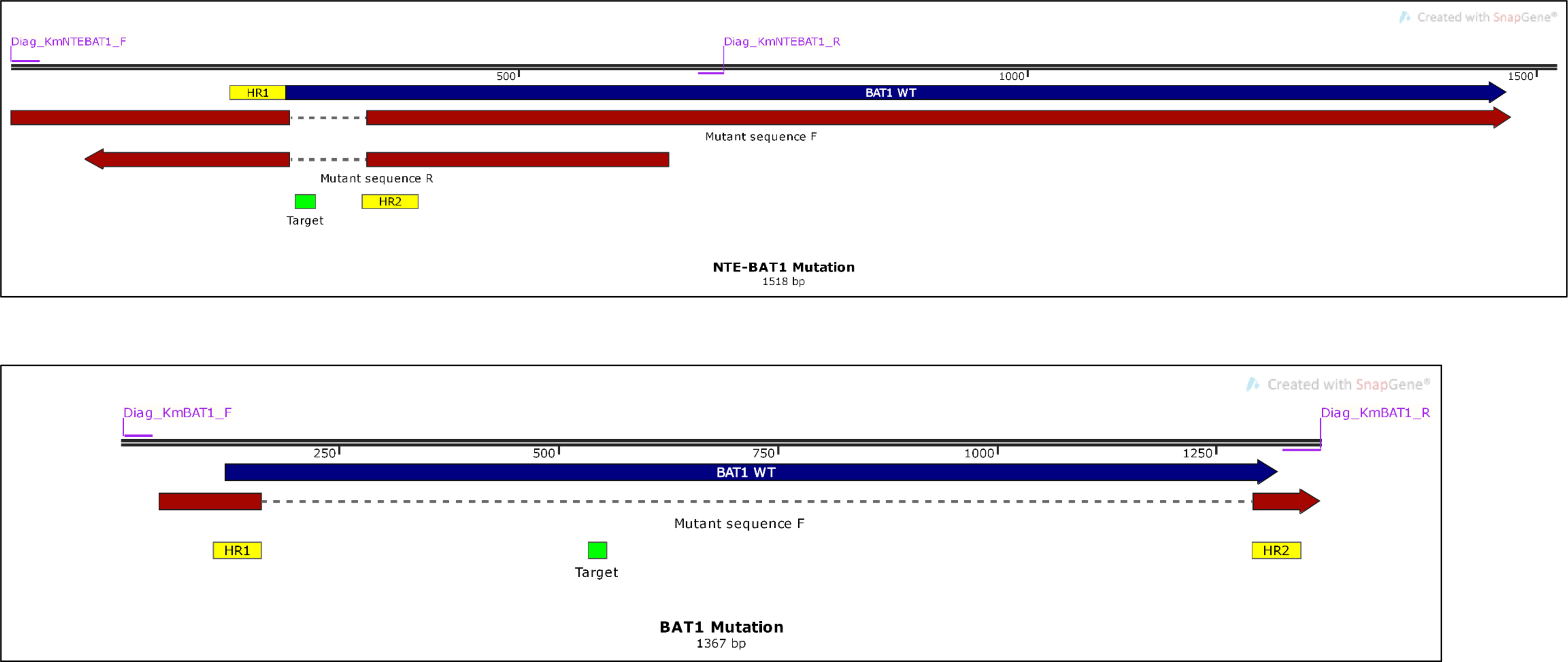

1 Gel electrophoresis of deletions:

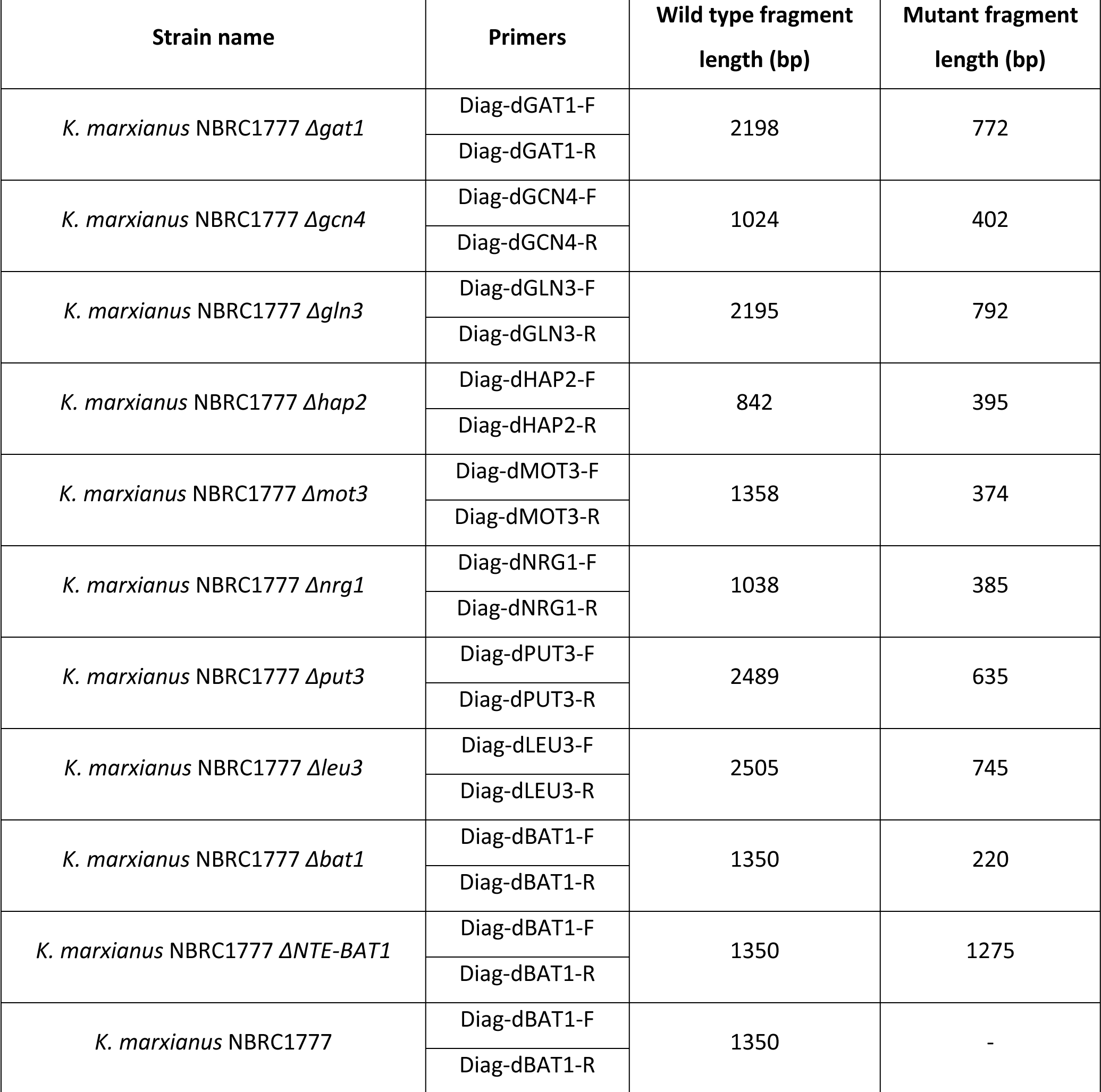
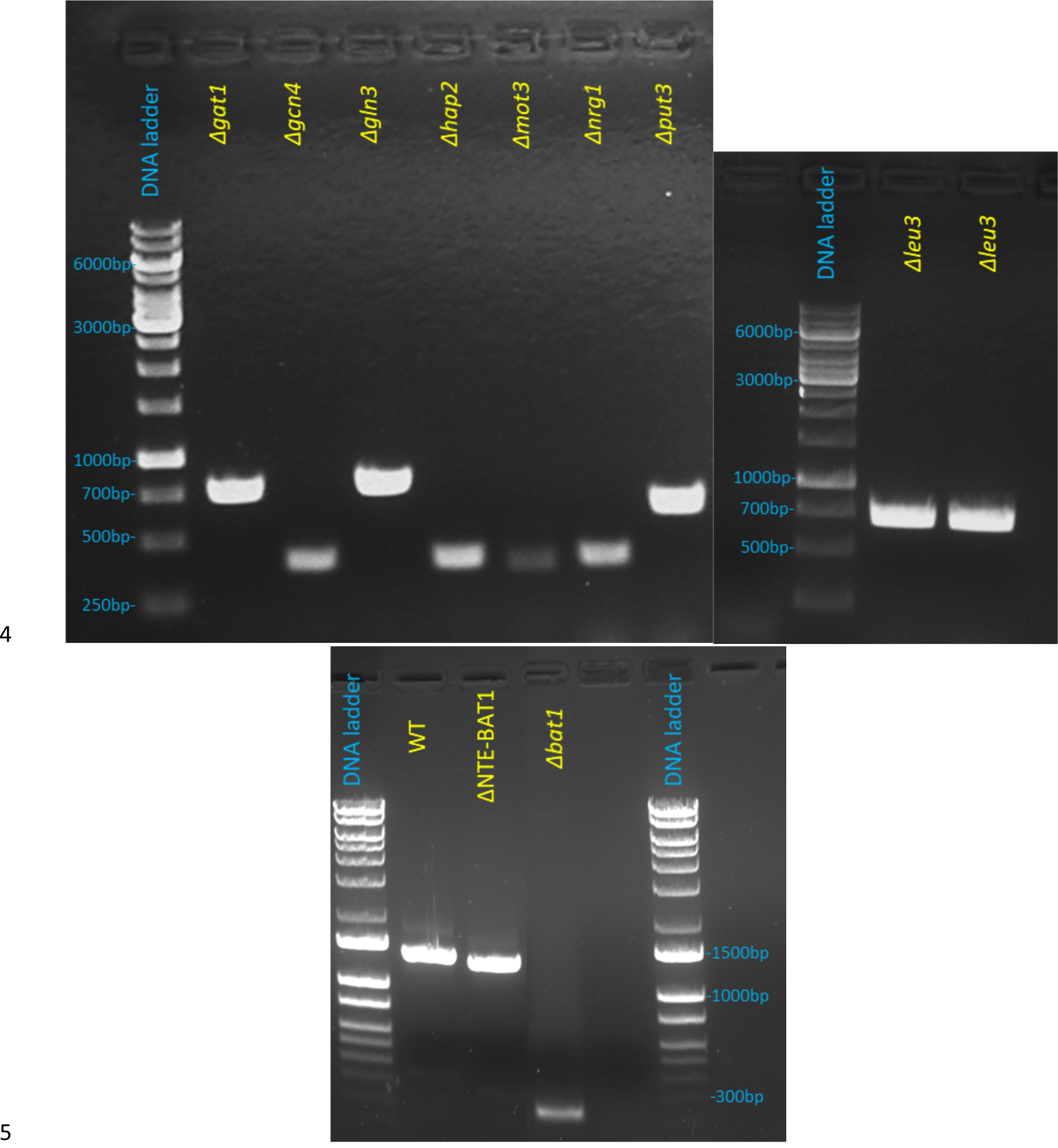

